# *Rhizoctonia theobromae* isolates causing Vascular-Streak Dieback of Cocoa and Cassava Witches’ Broom Disease are likely host-specific, regionally divergent and homothallic

**DOI:** 10.64898/2026.02.08.704712

**Authors:** Peri A. Tobias, Jacob M. Downs, Sylvie Nolf, Agus Purwantara, Junaid Muhammad, Eirene Brugman, Mareike Möller, Marcin Nowicki, Juan M. Pardo, David I. Guest

## Abstract

The fastidious basidiomycete *Rhizoctonia (Ceratobasidium) theobromae* is a biotrophic pathogen that causes Vascular-Streak Dieback (VSD) of *Theobroma cacao* (cocoa). The fungus has also been identified as the cause of an emergent disease known as Cassava Witches Broom Disease (CWBD) raising concerns that the pathogen is spreading to alternative hosts and to new regions. Interestingly, while VSD of cocoa and CWBD are reported as co-present in several countries, there is currently no evidence for cross-infection between species. The fungus is difficult to culture *in vitro* due its slow growth and Koch’s postulates have not been definitive on either host. The complete fungus life cycle therefore remains enigmatic, though studies have progressed knowledge on pathology within the both the cocoa and cassava hosts. We have conducted limited field trials and sequenced mating (MAT) and ITS loci of isolates from various infected hosts and regions. We hypothesize that (i) genetic variation at MAT loci correlates with region or host (ii) long amplicon ITS sequences between isolates are more definitive for polymorphisms (iii) life-cycle traits of *R. theobromae* may be inferred from MAT loci (iv) cassava grown under VSD infected cocoa will be infected and develop symptoms of CWBD. We did not find any cross-infection in field trials, and we show that the pathogen is highly homozygous, despite undergoing meiosis, indicating a predominantly homothallic life cycle. Our data indicate that the pathogen is likely host specific and regionally divergent and suggests that host specificity on cocoa and cassava evolved by selection from a common ancestor rather than a host jump.

## Introduction

The fastidious basidiomycete *Rhizoctonia theobromae* (P.H.B. Talbot & Keane) Samuels & Keane (syn*. Ceratobasidium theobromae*, *Oncobasidium theobromae, Thanatephorus theobromae*) is a biotrophic pathogen that causes Vascular-Streak Dieback (VSD) of *Theobroma cacao* L. (cocoa) (Keane et al., 1972). Within the *Rhizoctonia* binucleate anastomosis group, *R. theobromae* is related to the soil-borne pathogen, *R. solani* J.G. Kühn (Ajayi-Oyetunde and Bradley, 2018; Sperschneider et al., 2025) and to orchid mycorrhizae (Freestone et al., 2021; O’Donnell et al., 2025).

Once considered to be restricted to southeast Asia, a change in characteristic VSD symptoms on cocoa was noticed in Indonesia from 2004 onwards, from green-spotted leaf chlorosis to necrotic blotches, with both symptoms leading to tree death in susceptible hosts (Samuels et al., 2012). Molecular phylogeny, based on Internal Transcribed Spacer region (ITS), showed that there were three potential *R. theobromae* populations originating from Malaysia/Indonesia, Papua New Guinea and Vietnam (Samuels et al., 2012). The sequence variants did not correlate with changing symptoms but additional studies in Sulawesi, Indonesia, found a correlation with changing environmental conditions (Bryceson et al., 2023).

More recently, *R. theobromae* has been identified as the cause of an emergent disease known as Cassava Witches’ Broom Disease (CWBD) of *Manihot esculenta* Crantz simultaneously in Southeast Asia and the Americas (de Oliveira et al., 2025; Leiva et al., 2023; Pardo et al., 2024), and VSD disease of *Cercis canadensis* L. (Redbud) (Beckerman, 2022). These findings raise serious concerns that the pathogen, considered a new encounter pathogen of cocoa (Marelli et al., 2019), is spreading to alternative hosts and to new regions. Cassava is a staple food crop in tropical Asia, South America and Africa and is often grown within proximity to cocoa (Ntui et al., 2024). Should the pathogen be cross-infective and reach West Africa on cassava, also the major world producer and supplier of cocoa beans, the effect on both staple and economic crops would be devastating. Concerningly, recent sequence data from numerous CWBD isolates in the Philippines indicated that the pathogen haplotype on cassava is identical to the VSD pathogen on cocoa from Indonesia (Landicho et al., 2025) and validated earlier findings of regional variation (de Oliveira et al., 2025, Gil-Ordóñez et al., 2024). While the Philippines study results are interesting, the sequences investigated were limited to standard ITS and 28S ribosomal regions from diseased cassava only, and there is no field evidence that isolates from cocoa are virulent on cassava or other hosts, or whether the species includes host-specific populations or strains.

The fungus causing VSD on cocoa is difficult to culture *in vitro* due its slow growth, with colonies overrun by commensal fungi, thereby hampering inoculation studies. Short-lived basidiospores, released pre-dawn from rainfall-induced sporophores, are the assumed inoculum that infect young cocoa leaves (Keane and Prior, 1991). Koch’s postulates have not been definitive (Samuels et al., 2012), however basidiospores from sporophores have produced binucleate hyphae within xylem of cocoa leaves (Keane et al., 1972a; Keane and Prior, 1991). Similarly, Koch’s postulates are not confirmed for the pathogen causing CWBD, though limited culturing has been possible (Gil-Ordóñez et al., 2024; Landicho et al., 2025). The complete fungus life cycle and the disease cycle therefore remain enigmatic, though many studies have progressed knowledge on pathology within the cocoa and cassava hosts. Interestingly, while VSD of cocoa and CWBD are reported as co-present in several countries (Table 1), there is currently no evidence of cross-infection between species.

**Table 1.**
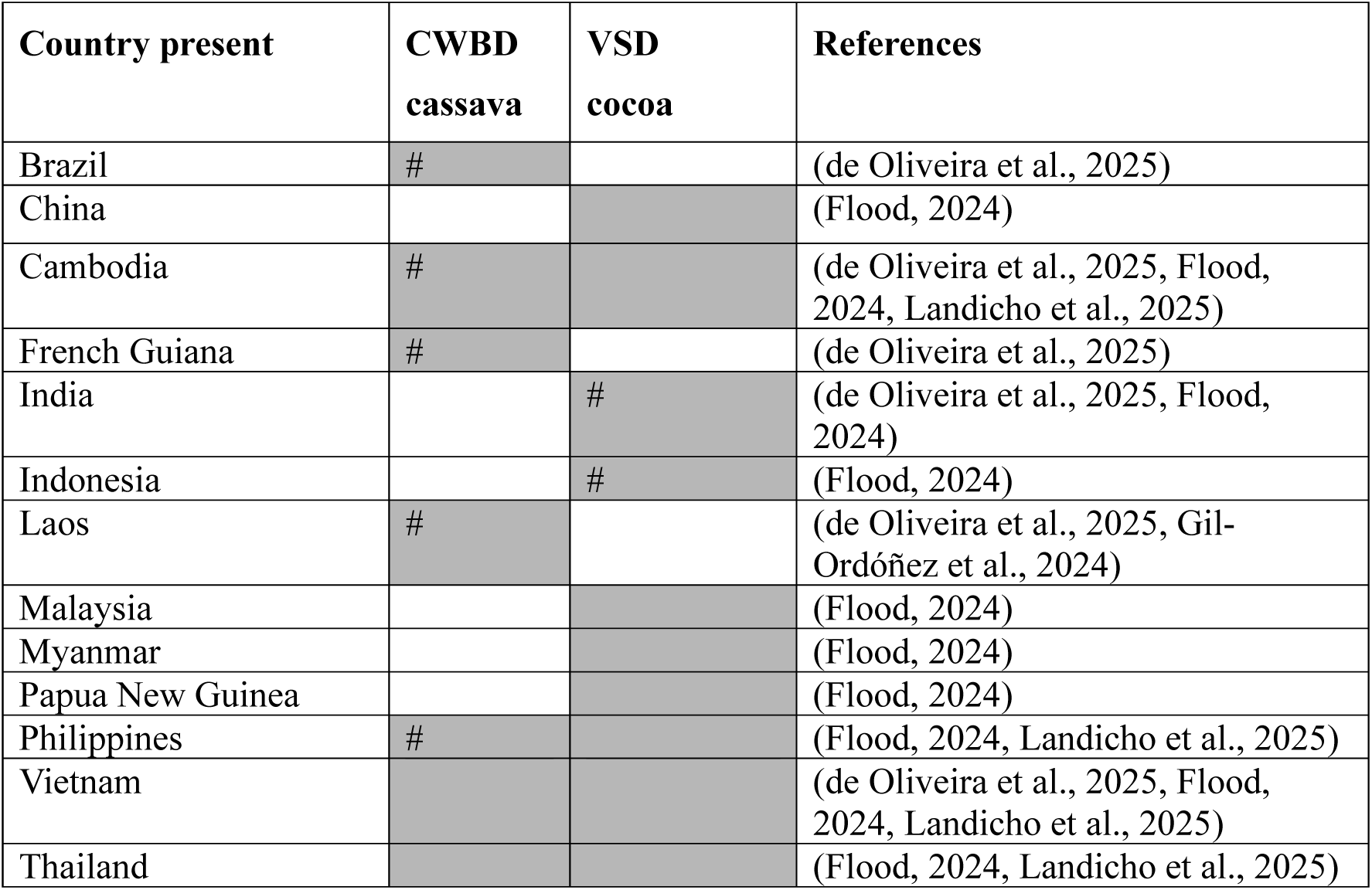
*Rhizoctonia theobromae* (syn. *Ceratobasidium theobromae*) reported (shaded cells) as causal agent of Vascular Streak Dieback of *Theobroma cacao* and of Cassava Witches Broom Disease on *Manihot esculent*a. # indicates molecular validation based on ITS.

All data currently used to identify *R. theobromae* isolates use short sequence amplicons derived from DNA regions from ITS and, more recently, Ca^2+^/calmodulin-dependent protein kinase (Leiva et al., 2023b; de Oliveira et al., 2025). The ITS regions are known to be good diagnostics for species but lack resolution to the sub-species or strain-level in fungi (Kauserud, 2023). Despite this lack of resolution, small variations are evident from the phylogenetic analyses of *R. theobromae* by geographic regions (de Oliveira et al., 2025) suggesting that these fungal isolates may be divergent at the strain level or higher. Additionally, the *Rhizoctonia* spp. clades have not previously been well resolved, potentially leading to misidentification in sequence databases (O’Donnell et al., 2025). The observed geographic variation in sequence data might help to explain the lack of reported cross-infection between distant host plant genera.

Fungal *homeodomain* (HD) transcription factors and *STE3-type **p**heromone **r**eceptors* (PR) at mating-type (MAT) loci are involved in sexual compatibility within the Basidiomycota (Bakkeren and Kronstad, 1994). Receptors encoded by the PR genes are biallelic in many basidiomycetes, meaning that each nucleus of the dikaryon has only one of the two divergent alleles (Luo et al., 2024), while a third PR gene is present as a conserved homolog in each nucleus (Ferrarezi et al., 2022), and is likely not involved in mate recognition. The HD loci are characterised by closely linked, bidirectional genes within each nucleus, referred to as bW1-HD1 (bW2-HD1) and bE1-HD2 (bE2-HD2) within some Pucciniales (Luo et al., 2024). The HD proteins from opposite and compatible nuclei heterodimerise to regulate mating transcription and are often polymorphic within the Basidiomycota, presumably leading to strain-level population diversity (Coelho et al., 2017).

Recent research, using phased genomes, determined strain-specific isolates for a rust-causing pathogen based on polymorphic HD loci, thereby providing a useful biosecurity tool (Ferrarezi et al., 2022; Luo et al., 2024). Though the HD loci within the closely related *R. solani* have not been found to be as polymorphic as many other basidiomycetes (Li et al., 2021), our preliminary investigation of putative HD genes within publicly available *R. theobromae* genomic resources (Ali et al., 2019; Gil-Ordóñez et al., 2024) shows sequence variation. The variation may therefore allow a useful approach to distinguish host and regional diversity within *R. theobromae* isolates from southeast Asia and elsewhere. Additionally, investigations of the MAT loci may help our understanding of the reproductive biology and the potential for new virulence to arise in *R. theobromae* if mating occurs between different strains.

Innovations in benchtop DNA sequencing permit full genes to be sequenced with Oxford Nanopore Technology (ONT), thus allowing greater depth of genetic differentiation. We used ONT MinION sequences from full gene amplicons, combined with bioinformatic approaches, to investigate genetic variation in *R. theobromae* isolates by region and host. Additionally, to investigate the potential for cross-infection from the pathogen causing VSD with CWBD, we conducted a field trial with cassava plants grown in proximity to heavily symptomatic *T. cacao* trees in Sulawesi, Indonesia.

Our research posed the following questions, (i) Does genetic variation at MAT loci (homebox-like, HD and *STE3*-type pheromone receptor genes) correlate with region or host and, if so, can these inform biosecurity management? (ii) Are long amplicon ITS sequences between isolates more definitive for polymorphisms? (iii) Can we infer *R. theobromae* life-cycle traits (homothallic versus heterothallic) from MAT-locus architecture and genome-wide heterozygosity? And (iv) do cassava plants develop CWBD under high natural inoculum exposure from adjacent cocoa trees with severe VSD symptoms. By integrating field exposure data, targeted long amplicon sequencing and whole-genome analysis, we aimed to delineate host range, population structure, and reproductive biology in this emergent pathogen complex.

## Results and Discussion

### No evidence for Rhizoctonia theobromae cross-infections from cocoa to cassava

There is no evidence that the pathogen causing CWBD on cassava and that causing VSD on cocoa can cross-infect. No data have been published demonstrating cross-infection despite disease symptoms on both hosts in several Southeast Asian and South American countries (Table 1). To our knowledge, there have been no reported symptoms of CWBD in Sulawesi where VSD is prevalent. To investigate the potential for cross-infection from the pathogen causing VSD with CWBD, we conducted a pilot field trial in South Sulawesi, Indonesia. Four individual cassava plants (Kahubi Palolo2 based on morphological traits) (Amin et al., 2024), two each sourced from Galonta and Lewaja in Enrekang Regency, were grown within immediate proximity of heavily VSD symptomatic *T. cacao* trees in South Sulawesi, Indonesia (Suppl. Table 1). *Rhizoctonia theobromae* sporophores were emergent on leaves of the VSD symptomatic trees (Experimental Procedures, Figure 7 C). After more than four months of field exposure over the rainy season (November – June in Enrekang Regency) no symptoms appeared on cassava. Leaf samples taken from the field-grown cassava and from the symptomatic cocoa trees, were surface sterilized and DNA was extracted. Using the ThanITS1/2 primer pair, that has been shown to be highly specific to *R. theobromae* (Samuels et al., 2012), we only produced expected 500 bp size amplicons from VSD symptomatic cocoa but not from exposed cassava (Suppl. Figure 1). These results show that, despite high exposure to pathogen propagules, there was no evidence of infection or latent colonization of the cassava leaves. While it is possible that the specific cassava genotype chosen for the field trial may be resistant to the pathogen, our results provide no evidence that *R. theobromae* from VSD-infected cocoa is pathogenic on cassava. This conclusion is further supported by sequence data described below.

### The Rhizoctonia theobromae genome is highly homozygous

We investigated the homozygosity and the predicted genome size for a Laos cassava *R. theobromae* isolate (LAO1; Gil-Ordóñez et al., 2024), using raw sequence data available at the National Center for Biotechnology Information (NCBI). Genome size calculations based on kmers indicated a diploid genome size of 87.5 Mb (43.7 Mb haploid). The published genome (LAO1) used binucleated cultured fungal hyphae and assembled to 33.4 Mb. We mapped the reads to the genome to investigate heterozygosity based on diploid Single Nucleotide Polymorphism (SNPs) and assessed the ploidy using GenomeScope (Ranallo-Benavidez et al., 2020). Prior to filtering the data for read depth and quality we found 8,283 heterozygous SNPs, however post filtering we found only 580 polymorphic nucleotides, indicating low genome-wide heterozygosity. Results based on 21 kmer histogram input to GenomeScope indicated very low heterozygosity at around 0.25 percent and repeat lengths at ∼26 percent of the haploid genome (Suppl. Figure 2).

The mapped reads were visually inspected around the predicted *homeobox domain transcription factor* (Hbox) and HD(E) genes using Integrated Genome Viewer (IGV) (Thorvaldsdóttir et al., 2013a), as these regions are known to be heterozygous and polymorphic in other basidiomycetes (Coelho et al., 2017). The specific gene loci inspected are homozygous within the LAO1 genome for *R. theobromae* (Figure 1). Interestingly, visualized mapped reads show a homozygous SNP adjacent to the predicted HDE region at 46 kb (Figure 1 B), indicating a likely assembly error, though read depth overall remains consistent for both loci (∼900–1000×). Despite this extreme genome-wide homozygosity, we identified biallelic and protein-level homozygous PR genes (see below).

**Figure 1.**
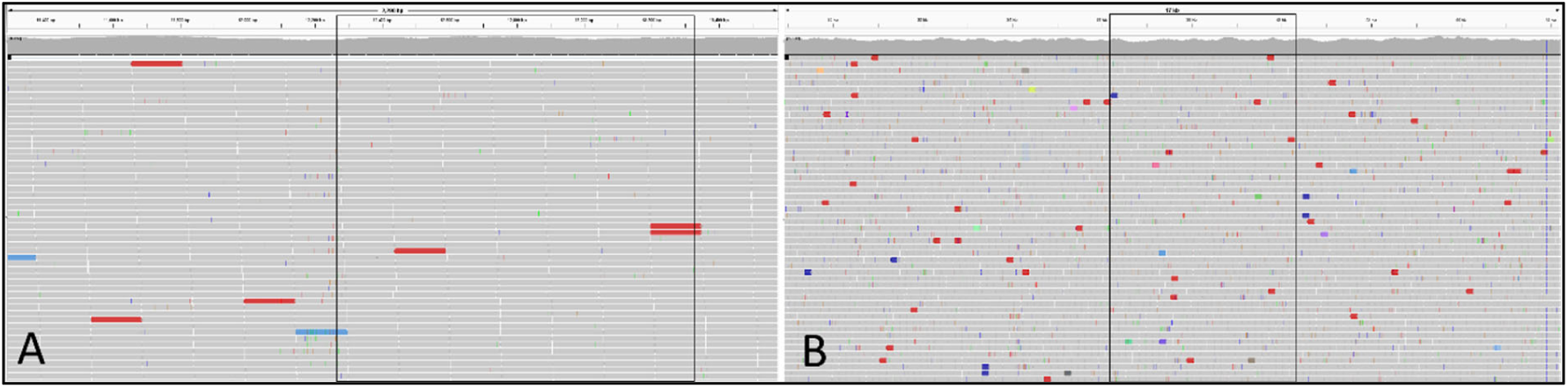
Trimmed Illumina reads mapped to *Rhizoctonia theobromae* LAO1 genome show high levels of homozygosity at the predicted *homeobox* (Hbox) (A) and HDE (B) loci visualised with IGV(Thorvaldsdóttir et al., 2013b). The boxed region identifies the predicted gene sequence region within the contig. DNA for the genome assembly was from cultured fungus isolated from symptomatic cassava with CWBD (Gil-Ordóñez et al., 2024a).

### Multiple pathogen isolates from various hosts and regions

Our detailed sequence analysis investigation used DNA extracted from 24 symptomatic host plants and from six fungal cultures (Table 2 and Suppl. Table 2). The isolates were selected to compare regions and host-specificity and to investigate MAT loci. Five culture samples were isolated from CWBD cassava from Laos and one from VSD symptomatic *Prunus persica* L. (Batsch), peach, from the USA. Symptomatic plant DNA included two from redbud and one from *Cornus mas* L., dogwood, from the USA, one from *Coffea arabica* L., a coffee plant in West Sulawesi, and 20 from cocoa trees growing in Sulawesi, Indonesia. These included historic isolate DNA available in our lab extracted from 12 symptomatic cocoa trees in Sulawesi (Junaid et al., 2021) and additional new leaf samples made from nine infected tree clones at the MARS Cocoa Research Institute, Pangkep, Indonesia in 2025. A total of 30 isolates were therefore used for comparative sequencing studies, although successful amplicons were not obtained from all isolates for all predicted loci. While we obtained Hbox sequences for all 30 isolates (Figure 3), our analysis compared a reduced set of nine isolates for all other genes investigated in this study (Table 2). The lack of amplicons for some genes from the cultured isolates from Laos and USA indicates likely sequence variation, as we obtained products from cocoa samples that were low quality mixed plant/pathogen DNA.

**Table 2.**
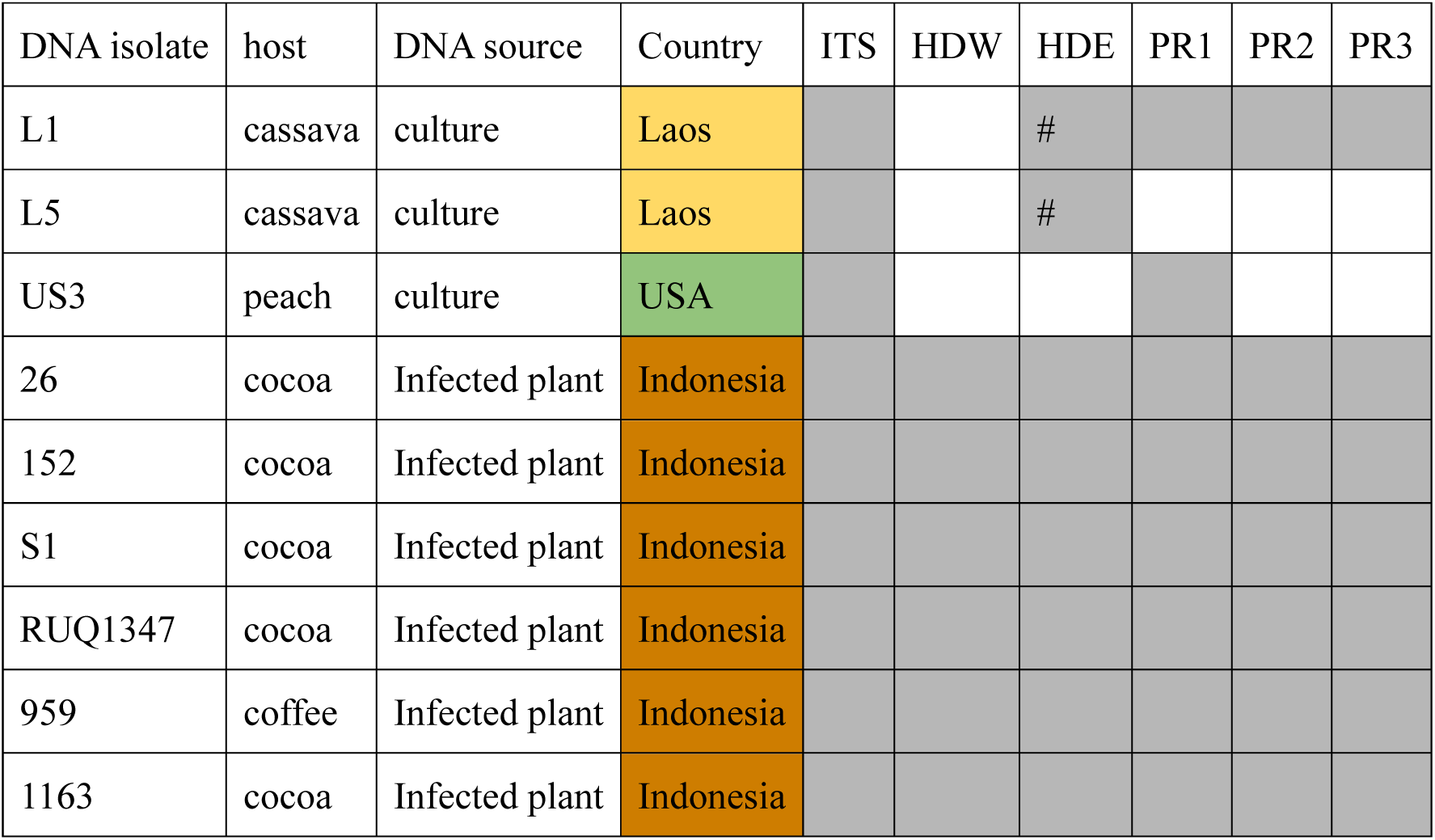
A reduced set of the *Rhizoctonia theobromae* isolates, the original host and successful complete gene amplicons sequenced from three geographic regions. ITS=internal transcribed spacer, and MAT loci predicted genes; HDW=homeodomain W, HDE=homeodomain E, PR1=*STE3* pheromone receptor 1, PR2= *STE3* pheromone receptor 2, PR3= *STE3* pheromone receptor 3. # indicates different primers for partial HDE (described in Experimental Procedures). Colours represent geographic locations; brown=Sulawesi, Indonesia, yellow=Laos, green=USA.

### ITS sequences are inconsistent for clarifying host-specific strains

To investigate the possibility of *R. theobromae* sequence variation for host specificity, we amplified long ITS sequences from eleven isolates, including eight from Sulawesi (cocoa and coffee), two from Laos (cassava) and one from USA (peach). We sequenced the 2 kb amplicons with MinION, mapped the reads to the full ITS sequence from the LAO1 genome and determined polymorphisms (Figure 2 B). To investigate the evolutionary relatedness, we included the predicted ITS region from a recently released Philippines *R. theobromae* genome (GCA_054508165.1) from CWBD. Notably, while the ITS from the coffee isolate aligned with all the cocoa isolates, one isolate from West Sulawesi (isolate 1163), aligned with alleles from Laos cassava and US peach isolates. The two cultured Laos CWBD isolates are heterozygous, indicated by designation A and B (Figure 2 A), and while isolate 1163 appears to be heterozygous, we cannot rule out that there may be two specific isolates within the mixed plant:pathogen DNA. All the other cocoa isolates appear to be homozygous for the ITS sequence. As we obtained many non-target reads from our sequencing, we decided to test whether isolate 959 was indeed from a coffee plant. We ran a blast search against the NCBI nucleotide database and confirmed that the sample was from *Coffea arabica* and not from *Theobroma cacao,* thereby indicating an additional host for the pathogen. Our long read ITS data shows that heterozygous isolates align in one clade and that homozygous isolates, apart from US3 from peach, align together. The data shows *R. theobromae* intraspecific strain variation, both within and between hosts, and indicates that the ITS locus can be inaccurate for determining fungal species or strains, as cautioned by Kauserud, (2023).

**Figure 2.**
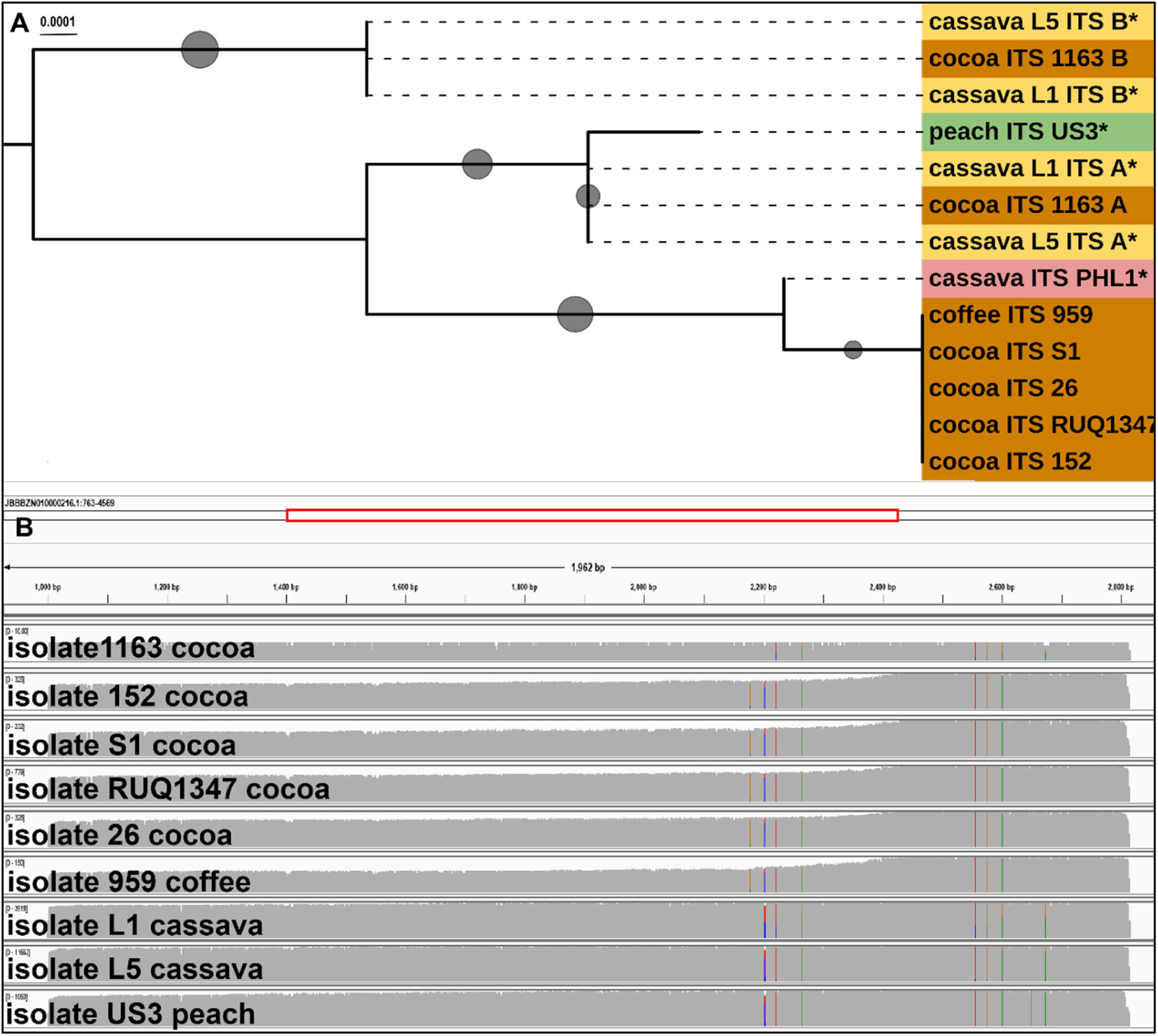
(A) Phylogenetic analysis of *Rhizoctonia theobromae* full (2 kb) ITS consensus sequences from Oxford Nanopore Technologies MinION amplicon reads and (B) single nucleotide polymorphisms visualised in IGV. The data shows homozygous Sulawesi cocoa isolates within one clade while the heterozygous cocoa isolate 1163, from West Sulawesi, aligns with Laos cassava isolates. Colours represent geographic locations and * indicates cultured isolate; brown=Sulawesi, yellow=Laos, green=USA, pink=Philippines. Where heterozygous alleles were identified they were designated A and B. Scale is nucleotide substitutions per site. The ITS region from a Philippines cassava pathogen genome (GCA_054508165.1) is included for comparison. The phylogeny was inferred using the Maximum Likelihood method and Tamura-Nei (1993) model of nucleotide substitutions. The Neighbor-Joining tree was generated using a matrix of pairwise distances computed using the Tamura-Nei (1993) model. Evolutionary analyses were conducted in MEGA12 (Kumar et al., 2024). Bootstrap values represented by grey circles range from 0.6-1.0. Tree midpoint rooted and visualised in iTOL (Letunic and Bork, 2021).

**Figure 3.**
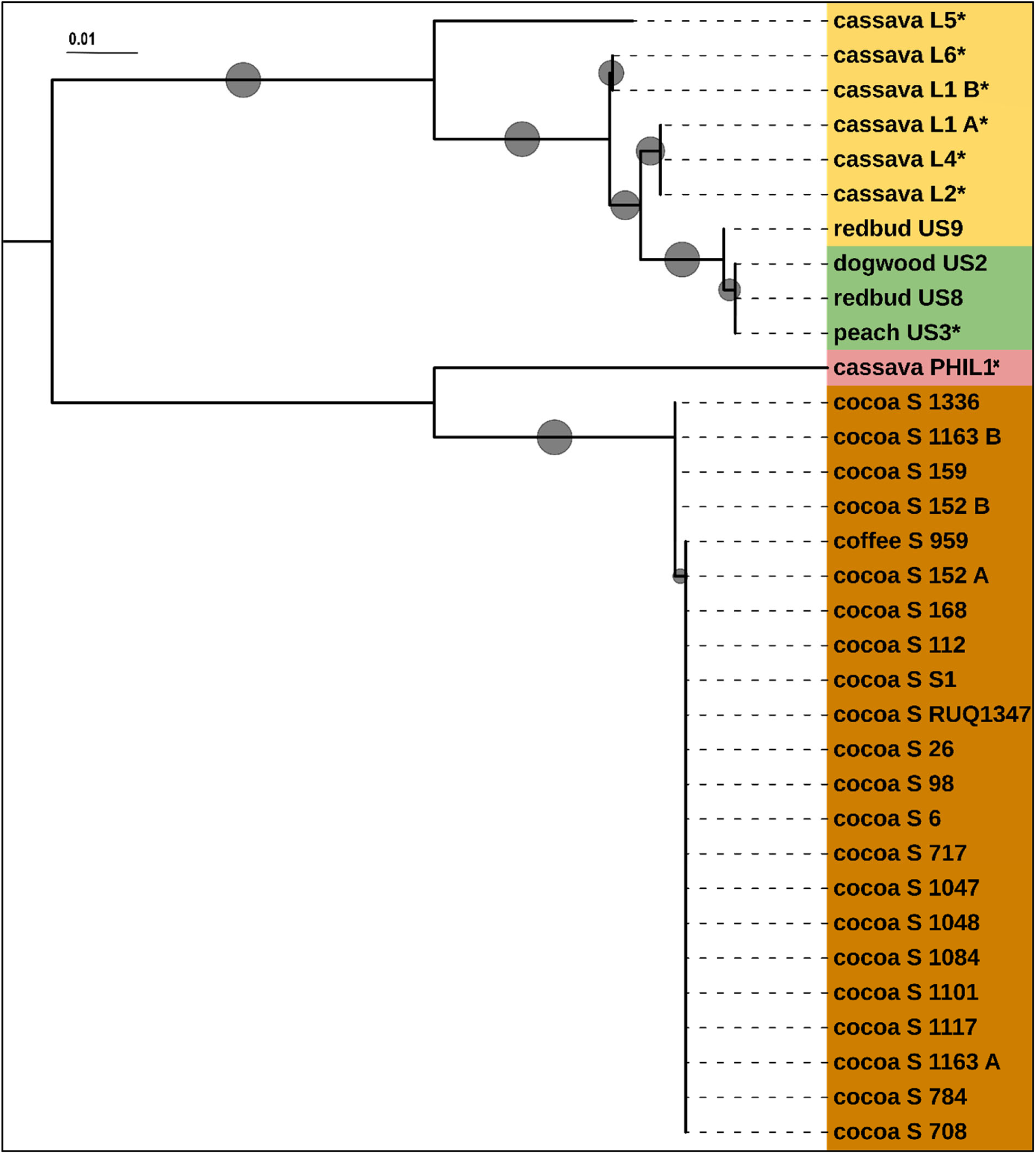
Alignment and evolutionary analyses were conducted in MEGA12 (Kumar et al., 2024) using Maximum Likelihood method with translated ONT sequenced amplicons from predicted *Rhizoctonia theobromae* homeobox (Hbox) genes. Isolates are from mixed plant:pathogen DNA unless represented by *=cultured isolate. Colours represent geographic locations; brown=Sulawesi, yellow=Laos, green=USA, pink=Philippines predicted sequence from the genome (GCA_054508165.1). Isolate 959 within the Sulawesi clade was isolated from a VSD symptomatic coffee plant. Heterozygous alleles designated A and B. Adaptive bootstrap results are indicated with grey circles by size (range: 0.6 – 1.0). Scale is amino acid substitutions per site. Tree midpoint rooted and visualised in iTOL (Letunic and Bork, 2021).

### Rhizoctonia theobromae homeobox predicted proteins are regionally and host distinct

We identified a putative *homeobox* (Hbox) gene sequence on contigs from publicly available *R. theobromae* genomes (Suppl. Table 3); Indonesian and Malaysian isolates infecting cocoa are named CT2 from Sulawesi (GCA_009078325) and Gudang 4, Tawau Sabah (GCA_012932095) respectively, and the Laos cassava isolate named LAO1 (GCA_037974915). We designed primers to amplify the full regions for sequencing. Although initial blast annotation suggested a homeodomain-like gene, domain architecture and genomic context indicate that this locus is not part of the canonical basidiomycete HD MAT locus; rather, it represents a *homeobox-like transcription factor* distinct from the typical paired HD genes. Nevertheless, our investigations show that this gene provides highly informative host– and region-specific haplotypes and therefore serves as a useful differential diagnostic for *R. theobromae* isolates. We used DNA extracted from regionally and host distinct isolates from 25 symptomatic plants and from six fungal cultures, described in the experimental procedures. We amplified, sequenced and annotated the complete Hbox genes across 30 *R. theobromae* isolates. Full Hbox sequences translated to predicted proteins (655 amino acid) with high regional and host homology (Figure 3).

Our data from Sulawesi cocoa isolates shows two alleles are present regionally. Sulawesi alleles differ by a single amino acid substitution (Alanine to Proline at 427 aa), whereas the Laos and US isolates, from varied hosts, grouped together into a separate clade (Figure 3). Notably the isolate from a VSD symptomatic coffee plant (isolate 959) from West Sulawesi clustered within the Sulawesi cocoa pathogen clade and isolate 1163 aligns within the cocoa isolates clade, despite the results for ITS described previously. Heterozygosity across all isolates is very low with only cocoa isolates 152 and 1163 and cassava isolate Laos 1 showing two alleles designated A and B (Figure 3). These data again indicate high levels of homozygosity likely a result of selfing.

The mixed plant:pathogen DNA isolates from 1163 and 152 could be interpreted as two distinct pathogens, or heterozygosity. However, the Laos L1 DNA was extracted from sub-cultured fungus and therefore it is more likely that the single isolate is heterozygous. Cultured *R. theobromae* isolates from cassava grown in Laos show three distinct morphospecies based on colony morphology and growth characteristics (pers. com. Juan M. Pardo), perhaps corresponding to these findings. Interestingly, we again included the predicted Hbox sequence from the Philippines *R. theobromae* genome (PHL1), cultured from diseased cassava, and found it more closely aligned with the pathogen on cocoa though in a separate clade (Figure 3).

Homeobox transcription factors are known to be important regulators of morphological differentiation and cellular function in fungi (Calvo et al., 2024), including sexual development, pathogenicity and response to environment. While our research inadvertently investigated the full sequences for this gene, as a homolog to the MAT loci HD genes, its structural arrangement may indicate a role beyond mating. Despite this, a combined evolutionary analysis indicates relatedness to the HDE and HDW predicted genes (Suppl. Figure 3). The notable sequence variation that we have identified across isolates makes Hbox a useful candidate in identifying regional and host specificity and confirms that using only ITS sequences may provide inaccurate results.

### Homeodomain (HD) MAT loci are distinctive by host and region

In Basidiomycetes, canonical HD MAT loci consist of two closely linked, divergently transcribed genes (often referred to as HD1/HD2 or BW/BE) whose protein products heterodimerize between compatible nuclei to regulate sexual development. The MAT loci HD genes are often polymorphic between strains within a species (Coelho et al., 2017), an adaptation likely to support outcrossing. We therefore investigated whether the *R. theobromae* pathogen isolates are polymorphic at HD loci. Though we had difficulty obtaining amplicons for the HD, with limited quantity and quality of DNA available, we successfully resolved these for a reduced set of isolates (Table 2) including from Indonesian cocoa isolates and some isolates from Laos. The lack of HDW amplicons from Laos isolates was not due to the quality or quantity of the DNA, as this was extracted from cultured samples. We did not locate a predicted HDW gene within the LAO1 genome and we were only able amplify the incomplete HDE sequence indicating sequence variants at these loci. We therefore consider the negative amplicon results informative and suggest that future studies investigate these loci in greater depth. Full gene products from the successful amplicons were sequenced and processed through our bioinformatic pipeline to compare across isolates and to identify heterozygosity.

Phylogenetic evolutionary relationships of HD amino acid sequences by clade show that the Sulawesi cocoa isolates group together (Figure 4). Only the isolates L1 from cassava, and 152 from cocoa, showed two alleles for HDE. Isolate 152 was the only isolate that indicated DNA heterozygosity (HDE A/HDW A and HDE B/HDW B), in the typical presentation for this locus. However, we found that, due to synonymous SNPs, the annotated proteins for HDE A and B were exact homologs unlike those predicted from *R. solani* (RsAG8-1) described in the experimental procedures. Our primers did not successfully amplify HD genes from the USA isolates from redbud and peach, suggesting sequence variations and potential validation that the *R. theobromae* isolates are host specific, as supported by our field trial. Together, the apparent high divergence of HDW in LAO1, failed amplification of HD loci from multiple Laos isolates and the protein-level homozygosity suggest that HD genes in these populations are unlikely to contribute substantially to mating-type diversification, even though they may still be required for sexual development *per se*.

**Figure 4.**
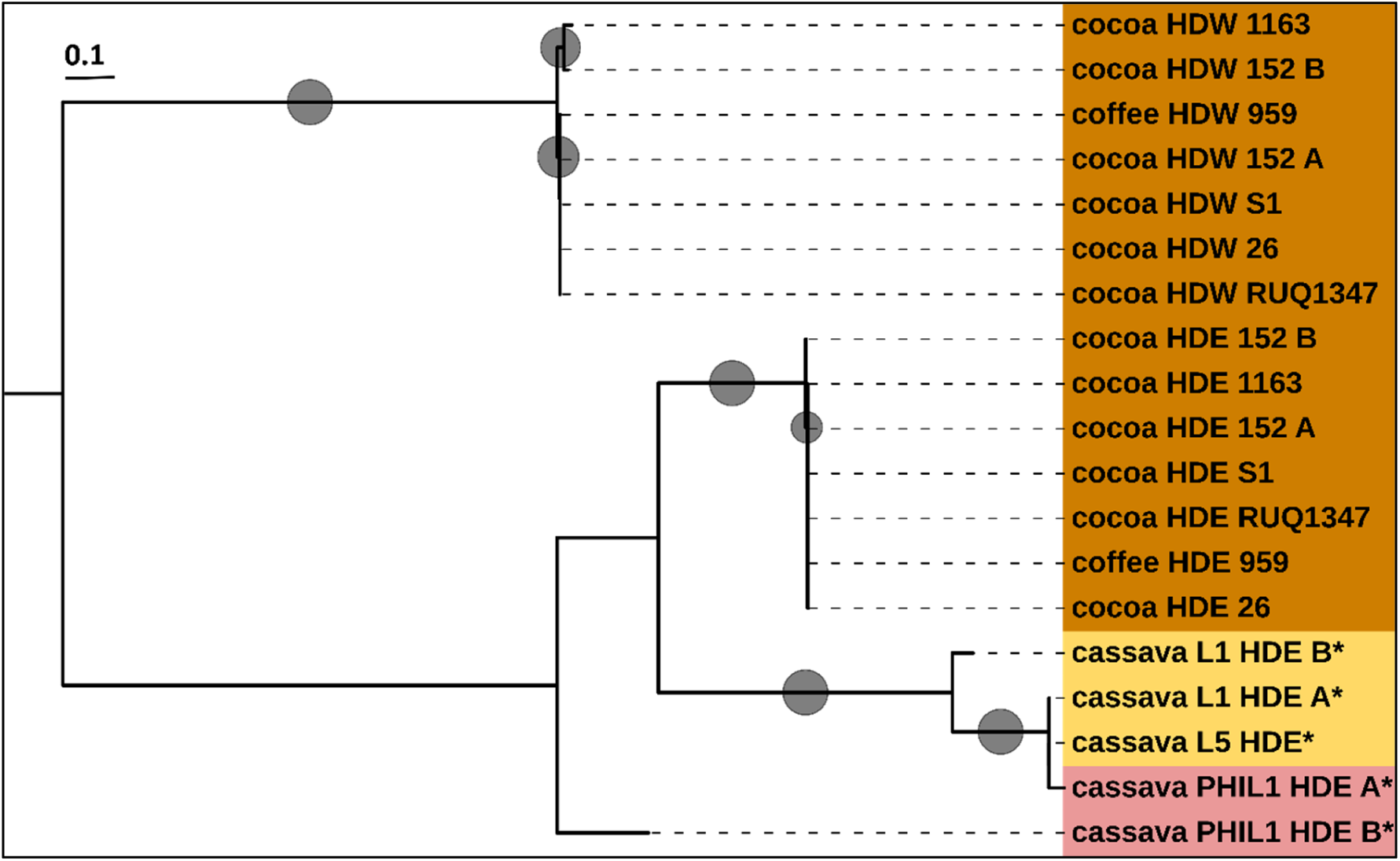
Alignment and evolutionary analyses were conducted in MEGA12 (Kumar et al., 2024) using Maximum Likelihood method with translated ONT sequenced amplicons from predicted *Rhizoctonia theobromae Homeodomain* west (HDW) and east (HDE) genes. Isolates are from mixed plant:pathogen DNA unless represented by *=cultured isolate. Colours represent geographic locations; brown=Sulawesi, yellow=Laos, pink=Philippines predicted sequence from the genome (GCA_054508165.1). Isolate 959 within the Sulawesi clade was isolated from a VSD symptomatic coffee plant. Heterozygous alleles designated A and B. Adaptive bootstrap results are indicated with grey circles by size (range: 0.6 – 1.0). Scale is amino acid substitutions per site. Tree midpoint rooted and visualised in iTOL (Letunic and Bork, 2021).

### MAT loci pheromone receptor (PR) amino acid sequences are closely related across regions and hosts

Basidiomycete MAT loci pheromone receptor genes are presumed biallelic and involved in recognition for plasmogamy (Coelho et al., 2017). We investigated *R. theobromae* pathogen isolates *STE3* pheromone receptor loci for these properties. Three predicted *STE3*-like Pheromone Receptor (PR) homologs, involved in fungal mating (Luo et al., 2024b), were found in all publicly available genomes for *R. theobromae*. We amplified and sequenced isolates and post-processed them to obtain haplotype alleles. Alignments using the translated consensus reads show three receptors by clade, with PR1 as likely *STE3.2-1* and PR2 and PR3 as *STE3.3* and *STE3.4* homologues (Figure 5). We were unable to amplify any PR candidates in Laos cassava isolates using our primers, but we successfully obtained amplicon data for PR1 in one US isolate from peach (Figure 5, US3). To compare our data with the publicly available *R. theobromae* genomes, Ct2 (Sulawesi), Tawau Sabah (Malaysia) both from cocoa VSD and LAO1 (Laos) and PHL1 (Philippines) from CWBD isolates, we extracted and translated the sequences for predicted PRs to include in alignment and phylogenetic analysis (Figure 5).

**Figure 5.**
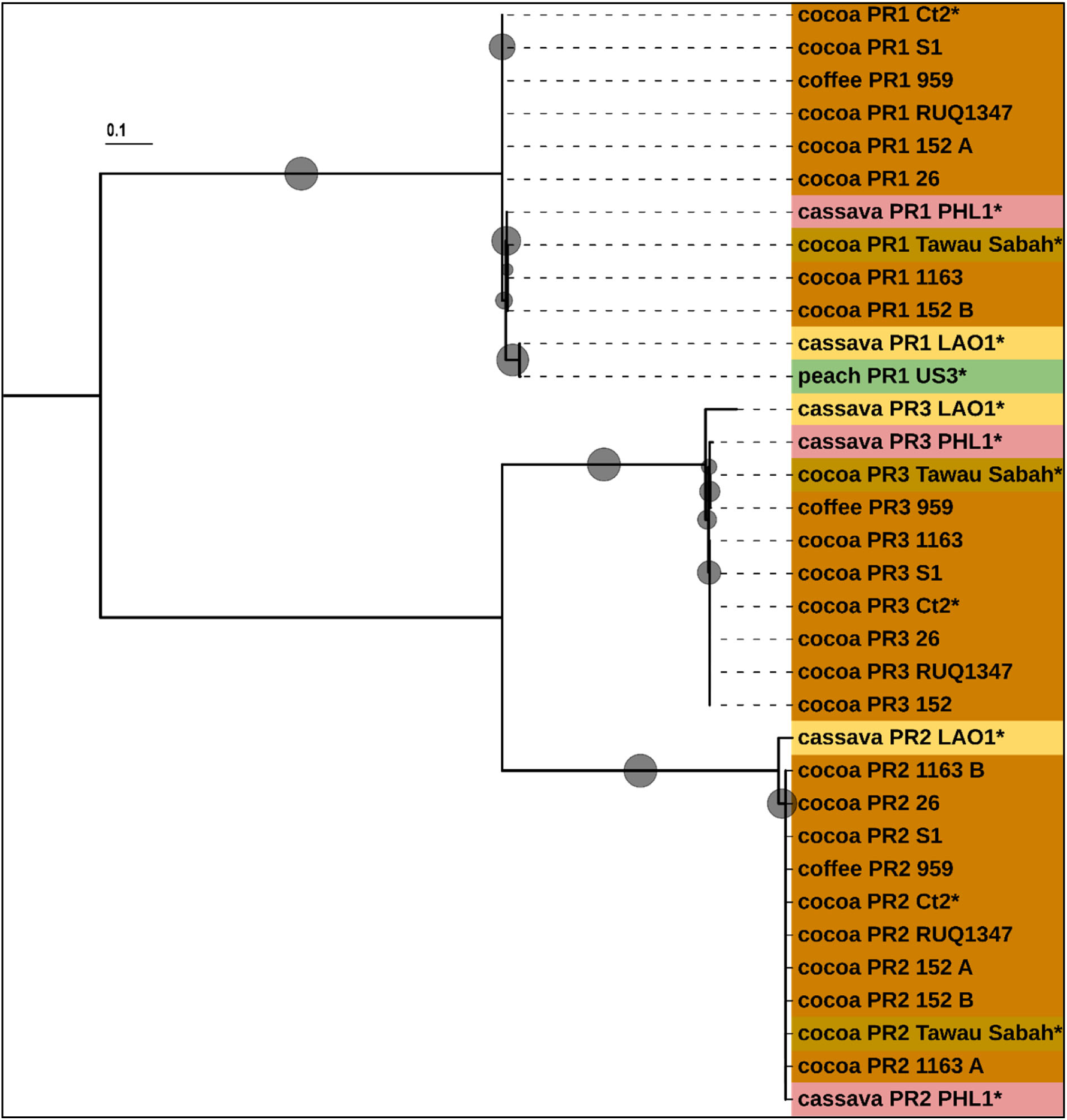
Alignment and evolutionary analyses were conducted in MEGA12 (Kumar et al., 2024) using Maximum Likelihood method with translated ONT sequenced amplicons from predicted *Rhizoctonia theobromae STE3 pheromone receptor* (PR) genes. Isolates are from mixed plant:pathogen DNA unless represented by *=cultured isolate. Colours represent geographic locations; brown=Sulawesi, light brown=Malaysia, yellow=Laos, green=USA, pink=Philippines. Predicted sequences from genomes are included for comparison and labelled PHL1=Philippine isolate (GCA_054508165.1), Ct2=Sulawesi isolate (GCA_009078325), LAO1=Laos isolate (GCA_037974915), and Tawau Sabah=Malaysia isolate (GCA_012932095). Isolate 959 within the Sulawesi clade was isolated from a VSD symptomatic coffee plant. Heterozygous alleles designated A and B. Adaptive bootstrap results are indicated with grey circles by size (range: 0.6 – 1.0). Scale is amino acid substitutions per site. Tree midpoint rooted and visualised in iTOL (Letunic and Bork, 2021).

Our data show that the translated PRs from the LAO1 cassava isolate are variant to all the cocoa isolates, and that PR1 from the US3 peach isolate aligns with the Laos isolate (Figure 5). We show that the cassava isolate PR sequences from the Philippines are evolutionarily very closely related to the cocoa VSD isolates from Indonesia and Malaysia, and that the predicted proteins are homologues. However, when we investigate the nucleotide alignment and evolutionary analyses of sub-trees for PHL1 PRs, we find divergence at the nucleotide-level (Suppl. Figure 4). Our data show all PR2 translated sequences from Sulawesi and Malaysia isolates are homologues, despite heterozygous DNA sequences for isolates 1163 and 152. Therefore, although we determined heterozygosity, synonymous SNPs show that these receptors are functionally homozygous. All PR3 alleles were shown to be homozygous, and our data show the isolate 959, found on coffee, shares an allele with the Tawau Sabah cocoa pathogen and is within the same clade as the PHL1 cassava pathogen.

The results for predicted PRs clearly separate heterozygosity at the DNA level from functional diversity at the protein level: whereas a small number of non-coding or synonymous variants are present, the encoded STE3-like receptors are essentially identical within host/region groups. Such protein-level homozygosity is consistent with a functionally homothallic system in which pheromone-receptor recognition specificity is tightly constrained, thereby limiting the potential for outcrossing between divergent mating types. It is interesting to note that predicted PRs within the *R. solani* genome assembly (Sperschneider et al., 2025), three homologues were present on chromosome 16A and B while only one was present on chromosome 12A and B, a pattern suggesting that PR2 and PR3 are tightly linked on chromosome 16. If a similar linkage occurs in *R. theobromae*, PR2 and PR3 would be inherited together on a single chromosome, thus allowing a germinating basidiospore to carry a complete, self-compatible set of pheromone receptors and thereby mechanistically enable homothallic fertility.

### Rhizoctonia theobromae isolates are predominantly homothallic

Evidence from genome-wide SNP analysis (LAO1), MAT*-*locus architecture (HD and PR genes), and published cytological studies of binucleate basidiospores collectively supports a predominantly homothallic reproductive system in *R. theobromae*. Based on LAO1 whole genome SNP analysis, and from investigation of specific alleles across multiple CWBD and VSD disease isolates, the fungus is predominantly homozygous, and likely selfing. Our results from other isolates also indicate that the homozygosity of the LAO1 isolate is not an anomaly. While the isolates we investigated are largely homozygous, some are heterozygous at single investigated loci including ITS, Hbox, PR1, PR2 and HD. These data suggest that there is potential for new differentiation to occur through mating or through somatic hybridization between strains, leading to new host compatibility. We note the symptoms in Indonesian cocoa leaves presented in the images (Table 2) indicate leaf apex necrosis where the pathogen is heterozygous for the alleles tested raising the hypothesis that rare recombinant or “heterothallic-like” isolates could contribute to the emergence of the novel symptom phenotypes noted in Indonesia since 2004. Testing this hypothesis will require targeted sampling of new symptom types, whole-genome sequencing to identify recombinant genomes, and, where feasible, controlled inoculations on cocoa and other hosts. (Bryceson et al., 2023b; McMahon and Purwantara, 2016; Parawansa et al., 2022)

It is well understood that *R. theobromae* hyphae are binucleate (Keane et al., 1972b). There is documented evidence that *R*. *theobromae* basidia produce four short-lived binucleate basidiospores that are known to germinate and penetrate the growing tip leaf (Keane et al., 1972b). Binucleate haploid basidiopores are well known for rust fungi and are presumed the result of mitosis (Anikster, 1983). The presence of basidiospores indicates a meiotic event and a full sexual life cycle (Keane, 2010; Keane and Prior, 1991b) despite plasmogamy and karyogamy not being detected to date. The full sexual stage generally requires both mating types (+ and –) however it has long been experimentally demonstrated that homothallic fungal mating arises from germinated haploid basidiospores (Buller, 1924; Buller, 1941). In cytological studies on *Agaricus bisporus*, three different meiotic types were identified that reproduced sexually, including homothallism (Kamzolkina et al., 2006). Previous work has shown microscopic evidence for *R. theobromae* basidiospores germinating and penetrating the leaf surface (Keane and Prior, 1991b). Our data appear to be consistent with investigations into STE3 pheromone receptor annotations from the phased *R. solani* genome (Sperschneider et al., 2025), where we identified paired homologues present in both nuclei. We suggest that the *R. theobromae* basidiospores have linked pheromone receptors, with PR2 and PR3 inherited on a single chromosome, as predicted for *R. solani*. Additionally, the observed homozygosity of HD genes in our datasets indicates that, in the isolates sampled, these loci are unlikely to contribute substantially to mating-type diversification, even though they may still be required for sexual development *per se*. The described arrangement would retain sexual compatibility within a single germinated spore, as represented in the *A. bisporus* study (Kamzolkina et al., 2006). This would indicate a homothallic and bipolar mating strategy (Wilson et al., 2015), partially explaining the observed MAT loci sequence results.

**Table 3.**
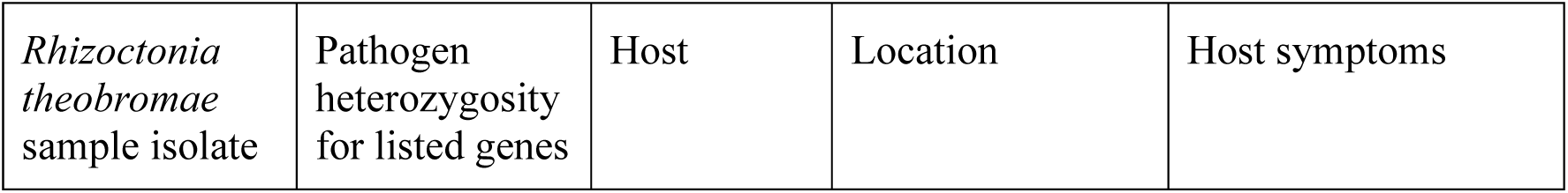
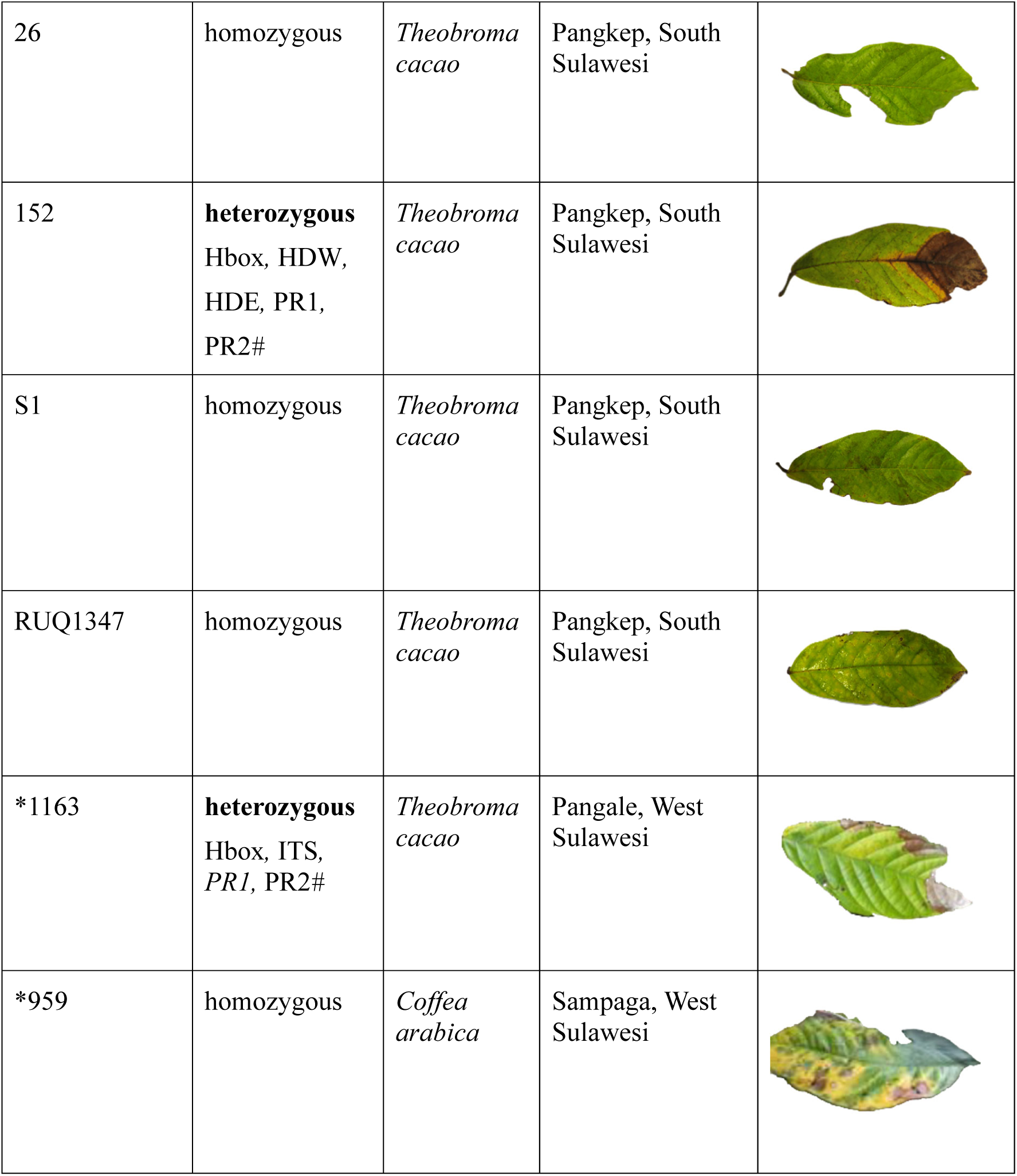
Sample subset of leaves used for plant/pathogen DNA extraction from current study. * Indicates historical DNA isolates used in the current study and associated images (Junaid, 2018). Pathogen heterozygosity of specific alleles investigated in the research. It should be noted that despite apparent heterozygosity for PR*2#* in two isolates, the amino acid translated sequences were identical. All images were edited to remove background with Microsoft Photos AI mode. Leaves are approximately 20 cm in length.

| *Rhizoctonia theobromae* sample isolate | Pathogen heterozygosity for listed genes | Host | Location | Host symptoms |
| --- | --- | --- | --- | --- |
| 26 | homozygous | *Theobroma cacao* | Pangkep, South Sulawesi |  |
| 152 | **heterozygous**<br>Hbox, HDW,<br>HDE, PR1,<br>PR2# | *Theobroma cacao* | Pangkep, South Sulawesi |  |
| S1 | homozygous | *Theobroma cacao* | Pangkep, South Sulawesi |  |
| RUQ1347 | homozygous | *Theobroma cacao* | Pangkep, South Sulawesi |  |
| *1163 | **heterozygous**<br>Hbox, ITS,<br>PR1, PR2# | *Theobroma cacao* | Pangale, West Sulawesi |  |
| *959 | homozygous | *Coffea arabica* | Sampaga, West Sulawesi |  |

### Synthesis of MAT locus structure and mating system for Rhizoctonia theobromae

Combining Hbox, HD and PR data presents a consistent picture. Hbox haplotypes provide clear host and region-associated signatures, whereas HD genes are highly conserved at the protein level and often apparently absent or non-amplifiable in certain lineages (for example, HDW in all cassava isolates). PR haplotypes are divergent by region only, with the PHL1 cassava isolate closely related for all PR predicted genes and variant to the LAO1 cassava isolate. These cassava PHL1 findings are taken from the NCBI publicly available genome and cannot be comprehensively compared with amplicon sequence data. However, the results point to a closer sequence relationship within the Southeast Asia region isolates from Malaysia, Indonesia and the Philippines. PR2 and PR3 are homozygous within host/region groups and, by analogy with *R. solani*, may be physically linked, thereby favouring self-compatible mating in a single nucleus. When integrated with the extremely low genome-wide heterozygosity estimated from LAO1, these features support a predominantly homothallic, clonally oriented reproductive strategy with occasional recombination events marked by localized heterozygosity at MAT*-*associated loci (Figure 6).

**Figure 6.**
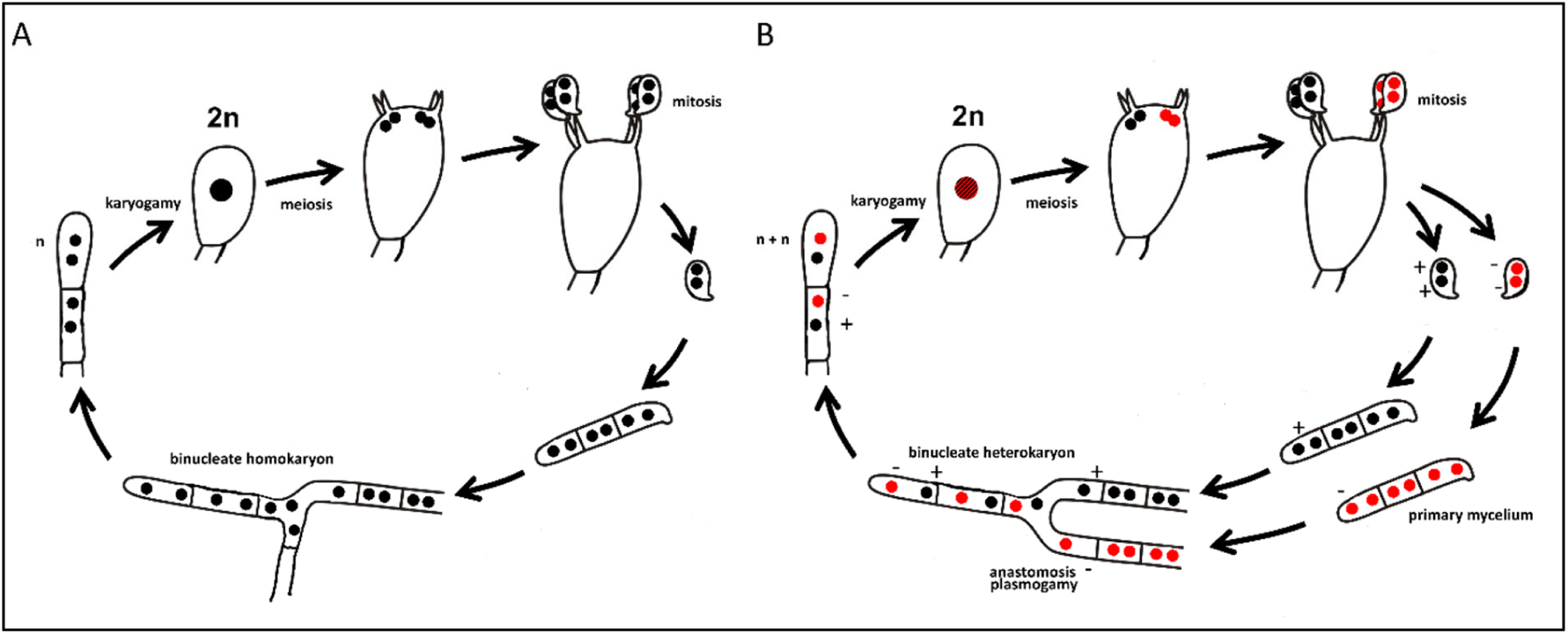
(A) Homothallic sexual reproduction for *R. theobromae*, maintaining a highly homozygous genome (B) and proposed heterothallic sexual reproduction with potential for new variation and new virulence. Life-cycle image adapted from M. Piepenbring© The heterothallic pathway and associated “new virulence” in panel B are presented as a conceptual model only and have not yet been demonstrated experimentally; they illustrate one plausible mechanism by which the limited heterozygosity observed at MAT associated loci could arise.

**Figure 7.**
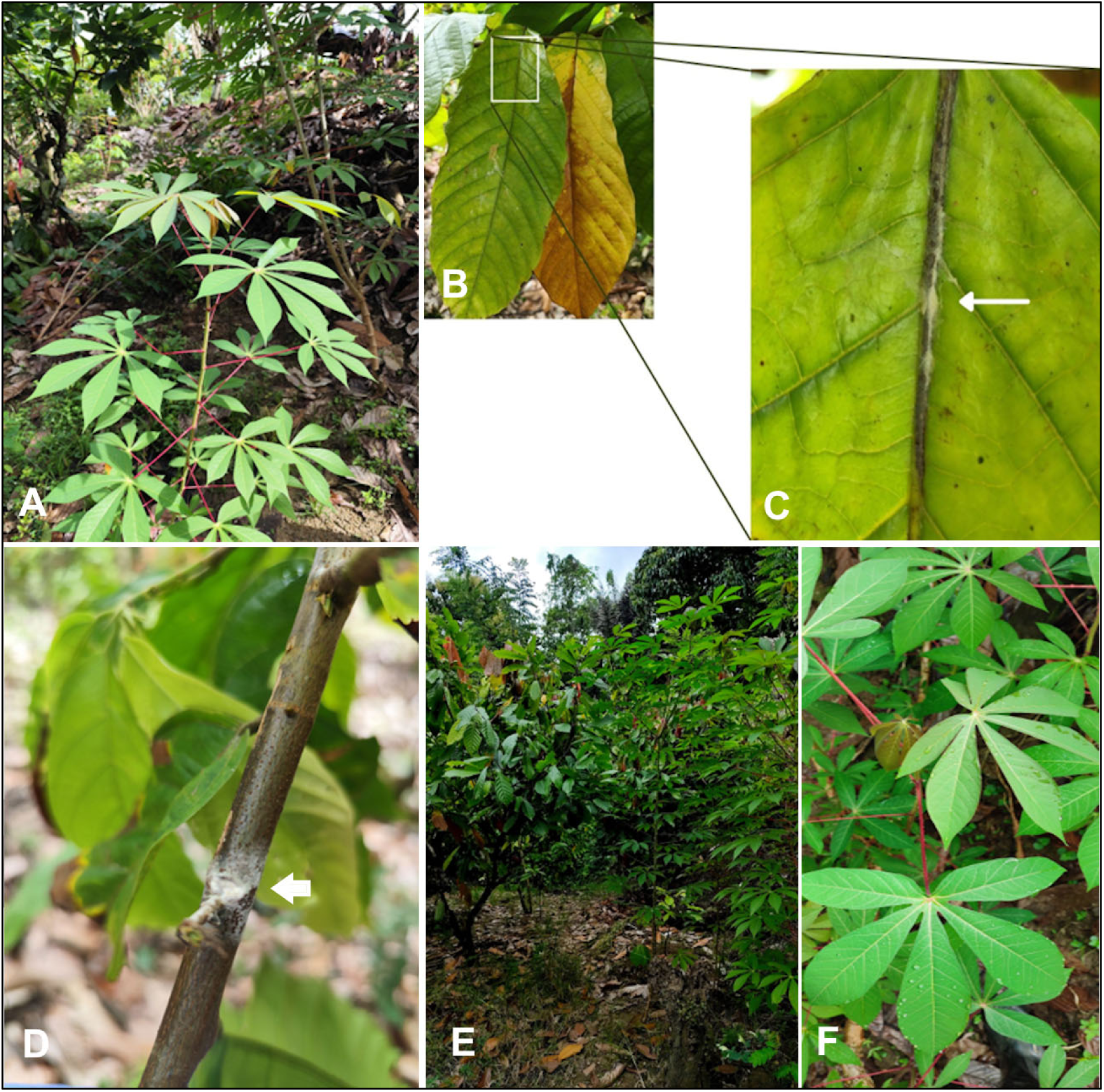
Healthy cassava plant (Palolo 2 accession) in Lewaja Field after two months growth surrounded by VSD pathogen infected *Theobroma cacao* trees. (A). *Rhizoctonia theobromae* pathogen sporophores emerging from cracking mid-vein (Arrow) (B and C) and from the leaf-scar (D) in the cassava/cocoa field trial. Healthy cassava plants (Palolo 2 accession) after three months growth surrounded by heavily VSD symptomatic and sporulating cocoa trees in Galonta Field (E and F).

## Concluding comments

Our results from both a field infection assay and sequence data point to regional and likely host-specificity for the pathogen causing VSD on cocoa. We determined no infection or disease latency on cassava grown under heavy exposure to *R theobromae* from VSD-symptomatic cocoa in a field trial conducted in Sulawesi, Indonesia. We confirm that ITS sequences are not consistent for determining strain-level *R*. *theobromae* diagnostics. Long amplicon ITS sequences reveal polymorphisms that do not consistently track with host or region. The sequence data for Hbox and HD genes is regionally and host specific however PR genes show conservation only by region. Our data support the hypothesis that there are multiple host-specific and regionally divergent variants of *R*. *theobromae* and suggest that host specificity evolved by selection from a common ancestor rather than a host jump from cocoa to cassava.

Whole-genome SNP analysis of the CWDB isolate LAO1, together with low heterozygosity at MAT-associated loci across multiple isolates, indicates an extremely homozygous, predominantly homothallic reproductive system, with occasional heterozygous loci (e.g. in ITS, HBOX, HD, or PR genes) suggesting rare recombinants. We propose that this reproductive mode favours long-term clonal persistence of host-adapted lineages, explaining the durable VSD resistance observed in cocoa crops. The pathogen likely retains a capacity for the emergence of novel variants when compatible strains co-occur. From a practical standpoint, our findings imply that cocoa and cassava co-cultivated in Southeast Asia are unlikely to share a single, freely cross-infective *R. theobromae* population, but that introduction of new lineages into naïve regions could still pose a major risk. The MAT-loci markers and primers developed here provide a toolkit for distinguishing host and region-associated lineages, monitoring pathogen movement and, in future work, testing hypotheses about the relationship between mating-type variation, symptom expression and host range.

Finally, our research data support the following proposed life-cycle; (i) The fungus is largely homothallic and can undergo full sexual cycle by selfing to produce haploid basidiospores that undergo mitosis, ie. two haploid nuclei, before release. (ii) Because it is selfing the pathogen is highly homozygous however our data shows two distinct and possibly linked pheromone receptor-types. Where isolates are heterozygous, plasmogamy and later karyogamy will introduce increased diversity and potential for new virulence (Figure 6 B). We suggest that future inoculation studies will benefit by understanding if the isolate is heterothallic or homothallic by prior testing with these primers. Additionally, if the germinated spores from two isolates are in proximity there is potential for plasmogamy leading to new variants.

## Experimental procedures

### Cross-inoculations and field trials

To investigate the potential for cross-infection from the pathogen causing VSD with CWBD, we conducted cassava field trials in Enrekang Regency, South Sulawesi, Indonesia. Four individual cassava plants, morphologically identified as accession Palolo2 (Amin et al., 2024), two each sourced from Galonta and Lewaja in Enrekang Regency, were grown within immediate proximity of heavily symptomatic *T. cacao* trees in Sulawesi, Indonesia. Cassava plants were planted in April (Galonta 1 and 2) and June (Lewaja 1 and 2) 2025 within 2-3 metres of a VSD infected *T. cacao* clone (data in Suppl. Table 1). The tree clone from a 20-year-old MCC02 rootstock with 15-year-old S2 entres (top-graft) showed symptoms of severe leaf necrosis, emerging sporophores from cracking petioles and branch dieback (Figure7). After 4-6 months of biweekly visual inspections, no symptoms on cassava were observed. To date, the plants continue to show no symptoms of VSD.

### DNA extraction and PCR amplification from cassava

To check for latent infection or endophytic presence of the pathogen within leaves of the cassava, approximately 3–5 g of the youngest cassava leaves (sprouts) were collected from each plant. The samples were wrapped in sterile tissue paper (Kimwipes), placed in zip-lock plastic bags, and stored in an ice cooler during transport. All samples were delivered to the laboratory and processed for DNA extraction within 24 h after collection.

Surface sterilization was performed by spraying the leaves with 70% ethanol and allowing them to air-dry for approximately 20–30 s. The leaves were then immersed in 1% sodium hypochlorite (NaOCl) solution for 30 s, followed by thorough rinsing with sterile distilled water (aquadest) to remove residual bleach. The sterilized leaves were gently swabbed dry using sterile tissue wipes. This procedure was modified from Alleyne et al. (2023). For DNA extraction, 100–200 mg of surface-sterilized leaf tissue was used for each sample. Genomic DNA was extracted using the Omega Bio-Tek HP Plant & Fungal DNA Kit (Omega Bio-Tek, USA) according to the manufacturer’s protocol. The extracted DNA was stored at −20 °C until further molecular analysis.

We ran PCR using the *R. theobromae* specific ThanITS primers 1 and 2 developed by Samuels *et al* (2012). These primers are species-specific for the pathogen, producing a DNA product at 500 bp. We included leaf samples from symptomatic *T. cacao* tree as a positive control for the pathogen. Previously we tested and validated the species-specificity of these primers on isolates of *Rhizoctonia* spp. isolated and cultured from several Australian orchid mycorrhizae including *Pterostylis saxicola, Genoplesium plumosum, Pterostylis mutica, Genoplesium superbum, Prasophyllum petilum, Pterostylis daintreana* and *Caladenia tessellate*. All were negative for *R. theobromae,* but positive using standard ITS1 and 4 primers (White et al., 1990). Additionally, we tested ThanITS primers on DNA extracted from asymptomatic cocoa plants, growing in the University of Sydney glasshouses, with negative results.

### Rhizoctonia theobromae DNA from culture and symptomatic plants

The sequence analysis used DNA extracted from 24 symptomatic plants and six fungal cultures. Five culture isolates were from CWBD cassava from Laos, isolated and DNA extracted according to the method described by Gil-Ordonez et al.(2024) and one from VSD symptomatic *Prunus persica* L. (Batsch), common name peach from USA. Symptomatic plant DNA included two from redbud and one from *Cornus mas* L., common name dogwood from USA, one from *Coffea arabica* L., a coffee plant in West Sulawesi, and 20 from cocoa growing in Sulawesi, Indonesia. Historic DNA isolates available in our lab were extracted from 12 symptomatic cocoa trees in Sulawesi (Junaid et al., 2021) and additional new leaf samples were made from nine infected tree clones at the MARS Cocoa Research Institute, Pangkep, Indonesia in 2025. DNA for new leaf samples was extracted using the Qiagen DNeasy Plant Pro Kit. A total of 30 isolates were therefore used for comparative sequencing studies. Details of isolates are available in Supplementary Table 2.

### Rhizoctonia theobromae homozygosity and genome size analysis

We accessed three publicly available *R. theobromae* genomes at NCBI; Indonesian and Malaysian isolates infecting cocoa named CT2 from Sulawesi (GCA_009078325) and Gudang 4, Tawau Sabah (GCA_012932095) respectively, and a Laos isolate infecting cassava, named LAO1 (GCA_037974915). Initially we investigated the homozygosity and the predicted genome size for LAO1 (Gil-Ordóñez et al., 2024) from cassava, as this was the only genome that had raw Illumina sequence data available at NCBI. We used raw fastq files with jellyfish (v2.2.6) (Marçais and Kingsford, 2011) and kmer setting 21mer. We mapped the reads trimmed with fastp (v0.19.6) (Chen et al., 2018) to the genome using hisat2 (v2.1.0) (Kim et al., 2019), and processed the output bam file to fix mate-pairs and deduplicate with samtools (v1.9)(Li et al., 2009).

To investigate heterozygosity, we visualised the kmer frequency histogram in RStudio (RStudio Team, 2015) and estimated diploid SNPs, using a bcftools (v1.22). The raw.vcf was filtered with bcftools using the following parameters: ‘QUAL<30 || DP<10 || DP>200’, shows 580 heterozygous SNPs, indicating low genome-wide heterozygosity. Additionally, we assessed heterozygosity by inputting the 21kmer histogram to GenomeScope (Ranallo-Benavidez et al., 2020) that indicated homozygosity at 99.7 percent and a haploid genome size around 81 Mbp. The predicted regions for the LAO1 genome Hbox and HDE loci are located on contigs JBBBZN010000017.1 and JBBBZN010000036.1 respectively (Figure 1 A and B).

### Identification of putative HD and PR genes for primer design

To identify putative HD genes, we used the three publicly available *R. theobromae* genomes and an isolate of *Ceratobasidium* sp. AG-Ba from China (GCA_016906575). All genomes are highly fragmented and haploid making investigations into MAT loci problematic. We first ran a local blast (Altschul et al., 1990) and a tblastn –evalue 10, with 773 annotated HD amino acid sequences derived from basidiomycetes downloaded from NCBI. The best hits against all genomes were found using Query= ELU37413.1 homeodomain transcription factor [*Rhizoctonia solani* AG-1 IA]. In this manner we identified a putative sequence on contigs from each of the genomes (Suppl. Table 3). We also ran a blastp search using ELU37413.1 with default parameters and the non-redundant protein sequence database (nr) on NCBI and determined a match with a predicted homeobox protein: KAB5589852.1 derived from the Sulawesi *R. theobromae* genome, at the same location and contig as determined with local blast. Based on information from other Basidiomycota HD allele arrangements (Luo et al., 2024b), we expected to find adjacent and closely linked HD genes (HDW and HDE) within each haploid nucleus of the *R. theobromae* however, we only found blast matches with a single gene. As the available genomes are all collapsed and fragmented haploid genomes, we assumed that the complete HD locus is present in the fungus and proceeded to design primers to amplify the complete region. We took the co-ordinates from these blast matches and extracted genomic regions 5 kilobases (kb) up and downstream using bedtools (v2.29.2) (Quinlan and Hall, 2010). We ran quick annotations on these regions using the on-line Augustus tool (v3.3.3) (Stanke et al., 2004), with *Ustilago maydis* as the reference. We then aligned the translated sequences from all genotypes to investigate homology. The amino acid sequences from the China isolate showed reduced homology and were excluded from further work. As we proceeded, the first chromosome-level and phased genomes for *Rhizoctonia solani* were made public (Sperschneider et al., 2025) permitting deeper investigation into the HD MAT locus. We ran blast of HD genes identified from *Ceratobasidium* Ag-Ba genome against the annotated proteins for AG8-1 and inspected the matches for co-location and orientation on the chromosome using IGV and gff3 files. In this manner, we determined two predicted HD genes that are opposite arranged and 295 bp apart on chromosome 11 (RsAG8-1_011449, RsAG8-1_011448 and RsAG8-1_026119, RsAG8-1_026120). These sequences were used in local blast against the three *R. theobromae* genomes, as before. Two predicted HD genes (here named HDW and HDE) were located for the Sulawesi and Malaysian genomes, on opposite strands and separated by ∼1,900 bp, but only HDE was found for the Laos genome. We proceeded to design primers based on the predicted HD locus, with the expectation that the previously located *homeobox* transcription factor gene (renamed for our study as Hbox) could be used as an additional differential between isolates. Similar methods were used to identify putative PR genes. We located putative PRs using blastp against *R. solani* predicted proteins with KAB5593758.1 Pheromone B beta 1 receptor [*Ceratobasidium theobromae*]. We identified predicted PRs from both haplotypes on chromosome 16A: RsAG8-1_014871, **RsAG8-1_014870**, RsAG8-1_014697, 16B **RsAG8-1_029556**, RsAG8-1_029557, **RsAG8-1_029374** and chromosome 12A RsAG8-1_011877 12B: RsAG8-1_026608. These data might indicate alternative mating mechanisms than identified across other basidiomycota, or that only those with predicted complete STE3 domains, noted here in bold, are functional. For *R. theobromae*, homologs from each of the genomes (Sulawesi, Malaysian and Laos isolates) of amino acid sequence translations were named PR1, PR2 and PR3. Sequence divergence between *R. solani* and *R. theobromae* made comparative analysis difficult but based on our findings we developed primers to try to resolve these sequences for isolates from different hosts and regions.

### Primers and conditions for long amplicon polymerase chain reaction (PCR)

Primers for the Hbox gene were designed using Primer-Blast (NCBI, 2024) with parameters set to DB: RefSeq, organism=Ceratobasidiaceae, max amplicon=4000 and all other defaults. Primer pair dimers were checked in-silico (Thermofisher Scientific, 2024) and locations visually examined in IGV against the three *R. theobromae* genomes using the motif tool. Primer pairs were selected to span the predicted full coding region of the Hbox gene. Other primers were designed using alignments in Geneious® and SnapGene® software. Long ITS primers were designed and generously provided by Dr Abigail Graetz from the Australian National University. All successful primers used in the current study are listed in Table 2 with the following conditions: 50 µL reactions using PrimeStar GXL® or Q5 High-Fidelity® DNA Polymerase, with 0.25 mL BSA (50 mg/mL) and annealing temperatures Hbox 60 °C, HDW/E 65 °C, PR1 58 °C, PR2 60 °C, PR3 62 °C, ITS 60 °C.

**Table 4.**
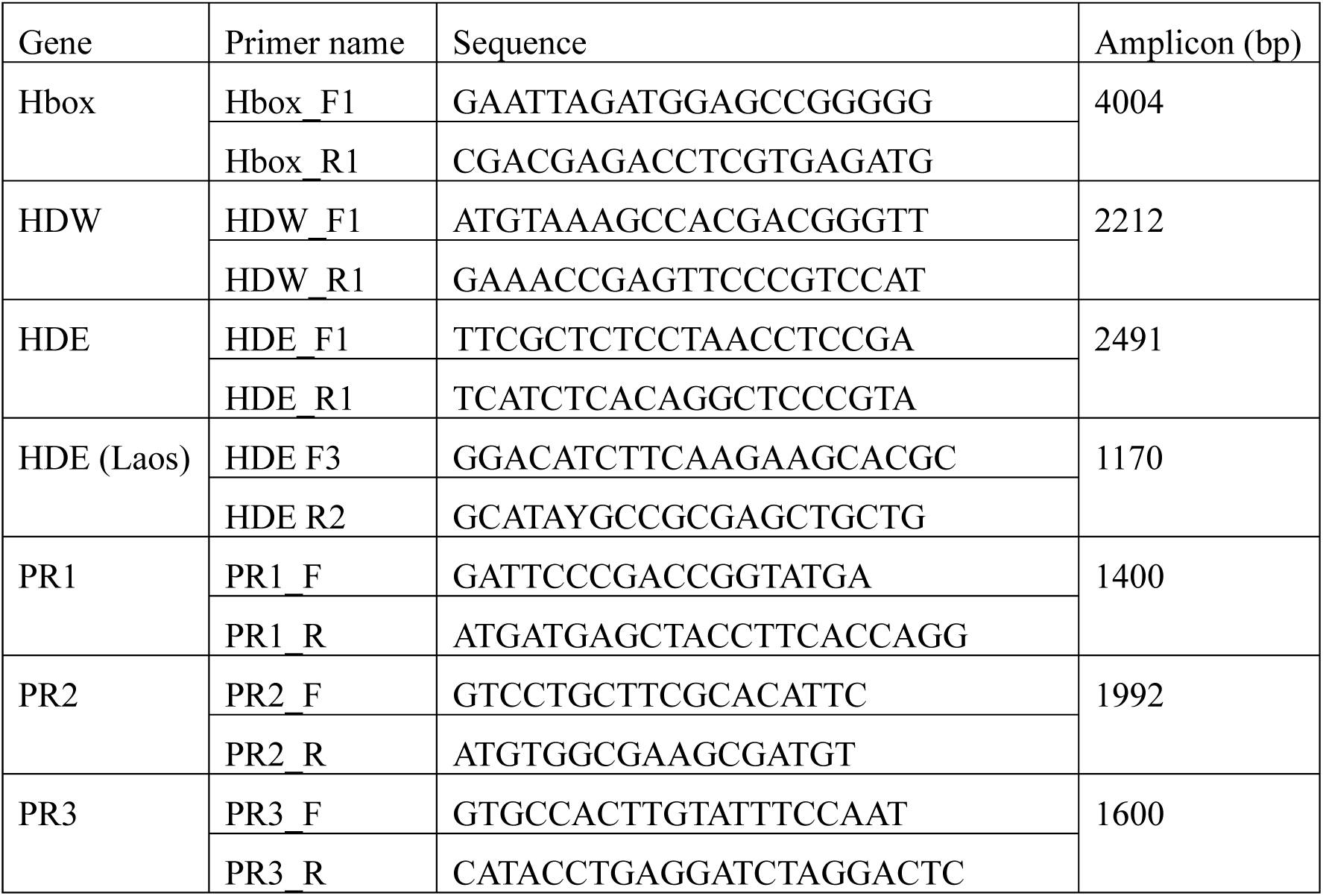
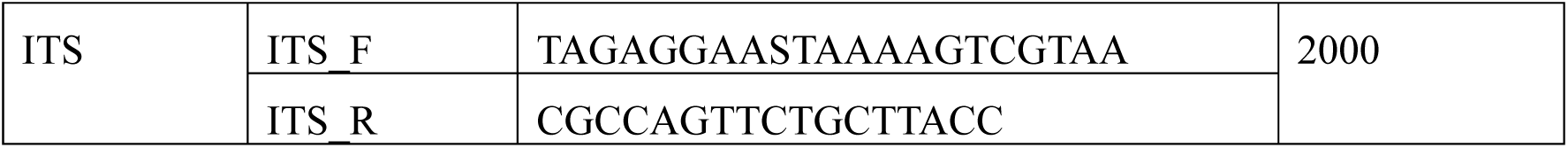
Primers designed to span full genes, except for HDE (Laos), for ONT MinION long amplicon sequencing from *Rhizoctonia theobromae*.

### DNA Library preparation and sequencing on MinION

Oxford Nanopore Technologies (ONT) Native Barcoding Kit 24 V14 and protocol were used. Sequencing runs were set up on R10.4.1 flow cell on a MinION Mk1B/D. Minimum read length set to 1 kb and minimum Q score=9. Basecalling was performed using Dorado 7.9.8 and the High-accuracy – v5.0.0 – 400bps model.

### Haplophasing and building consensus sequences from MinION reads

Our work necessitated an in-house pipeline to clearly identify heterozygous alleles and to produce allelic consensus reads for annotation and analysis. While ONT tools such as Epi2me (Epi2me Labs, 2024) are useful for homozygous reads, they are not suitable for heterozygous data. Details of the post-processing pipeline are available at https://github.com/alienshe/cocoa_project_scripts/tree/main. In brief, barcoded raw sequence reads were trimmed using porechop (v0.2.4) (Wick and Volkening, 2018). The trimmed reads were then aligned against the appropriate reference sequence using the map-ont function from minimap2 (v2.28-r1209) (Li, 2018). We used sequences spanning the gene of interest as the reference for mapping the trimmed reads rather than the full genome. From these alignments, a phased vcf file identifying variant SNPs and indels was made using Clair3 (v1.0.11) with the following arguments: platform=“ont”,include_all_ctgs, chunk_size=5000, enable_variant_calling_at_sequence_head_and_tail, enable_phasing, var_pct_full=1.0, ref_pct_full=1.0, and using the “r1041_e82_400bps_hac_v500” model (Zheng et al., 2022). We then used devider (Shaw et al., 2025) to identify haplotypes from identified SNPs. Additionally, the haplotag function from WhatsHap (v2.8) was used to sort reads into haplotypes (Martin et al., 2016) based on SNPs and indels. Finally, consensus sequences were made for each haplotype using the samtools (v1.21) consensus function (Li et al., 2009). For the ITS reads, where multiple non-target reads were detected, we filtered for sequences that included a match with ThanITS primers using vsearch (v2.21.1) and the following parameters; id_threshold=0.98 and cov_threshold=0.9. Output reads were then processed with our haplophase bash script, visualised with IGV (v2.13.0) and aligned with ClustalW (Thompson et al., 1994) to predict the evolutionary relationships with the Maximum Likelihood method within Mega12 (Kumar et al., 2024) graphic user tool. For Hbox and HD sequences we predicted annotations from consensus sequences using the Augustus web server and aligned as described previously. Predicted open reading frames for PR sequences were determined with the Emboss Transeq server (https://www.ebi.ac.uk/jdispatcher/st/emboss_transeq) and aligned as described previously within Mega12.

## Acknowledgements

Abigail Graetz from ANU designed and kindly shared the long amplicon ITS primers.

## CRediT author contributions Data availability

The data that support the findings of this study are available from the corresponding author upon request.

## Indonesian Foreign Research Permits (BRIN)

Peri Tobias 875/SIP/IV/FR/12/2025

David Guest 876/SIP/IV/FR/12/2025

Jacob Downs 877/SIP/IV/FR/12/2025

## Supporting Information Legends

**Suppl. Figure 1. Gel electrophoresis results of PCR amplified *Rhizoctonia theobromae* species-specific Than_ITS1/2 from cassava and VSD infected cocoa**

**Suppl. Figure 2. Linear GenomeScope (Ranallo-Benavidez et al., 2020) plot for 21kmer analysis of mapped Illumina reads to the *Rhizoctonia theobromae* LAO1 genome showing low heterozygosity and high repeat content.**

**Suppl. Figure 3. Alignment and evolutionary analyses were conducted in MEGA12 (Kumar et al., 2024) using Maximum Likelihood method with translated ONT consensus sequences from predicted *Rhizoctonia theobromae* homeobox (Hbox) and HD genes. Isolates are from mixed plant:pathogen DNA unless represented by *=cultured isolate. Colours represent geographic locations and * indicates cultured isolate; brown=Sulawesi, yellow=Laos, green=USA, pink=Philippines predicted sequence from the genome (GCA_054508165.1). Isolate 959 within the Sulawesi clade was isolated from a VSD symptomatic coffee plant. Heterozygous alleles designated A and B. Adaptive bootstrap results are indicated with grey circles by size (range: 0.6 – 1.0). Scale is amino acid substitutions per site. Tree midpoint rooted and visualised in iTOL (Letunic and Bork, 2021).**

**Suppl. Figure 4. Subtree alignment and evolutionary analyses were conducted in MEGA12 (Kumar et al., 2024) using Maximum Likelihood method with consensus ONT nucleotide sequences from predicted *Rhizoctonia theobromae STE3 pheromone receptor* (**PR**) genes. Isolates are from mixed plant:pathogen DNA unless represented by *=cultured isolate. pink=Philippines. Predicted sequences from genomes are included for comparison and labelled PHL1=Philippine isolate (GCA_054508165.1), Ct2=Sulawesi isolate (GCA_009078325), LAO1=Laos isolate (GCA_037974915), and Tawau Sabah=Malaysia isolate (GCA_012932095). Isolate 959 within the Sulawesi clade was isolated from a VSD symptomatic coffee plant. Heterozygous alleles designated A and B. Adaptive bootstrap results are indicated with grey circles by size (range: 0.6 – 1.0). Scale is nucleotide substitutions per site. Tree midpoint rooted and visualised in iTOL (Letunic and Bork, 2021).**

**Suppl. Table. (sheet one) *Theobroma cacao/ Manihot esculenta* cross inoculation field trial data to determin susceptibility to VSD in cassava. (sheet two) *Rhizoctonia theobromae* DNA isolates obtained and used in the current study from culture and symptomatic plants. (sheet three) Results from a tblastn search against three publicly available genomes for *Rhizoctonia theobromae* and one *Ceratobasidium* sp. for gene homologs investigated in the current study.**

