## Supplementary figures and images for "*Rhizoctonia theobromae* isolates causing Vascular-Streak Dieback of Cocoa and Cassava Witches’ Broom Disease are likely host-specific, regionally divergent and homothallic"

### Suppl. Figure 1.

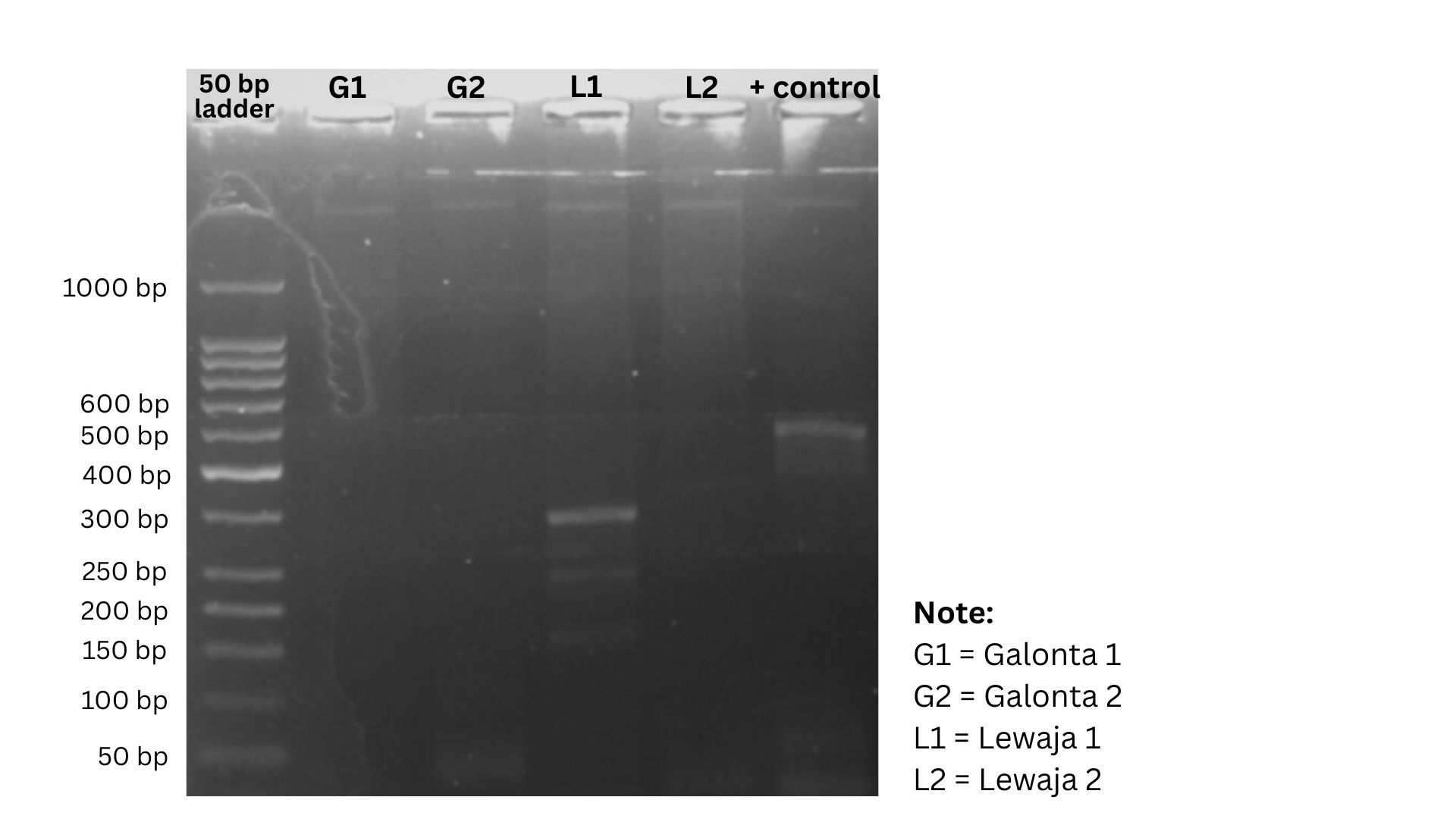

### Suppl. Figure 2.

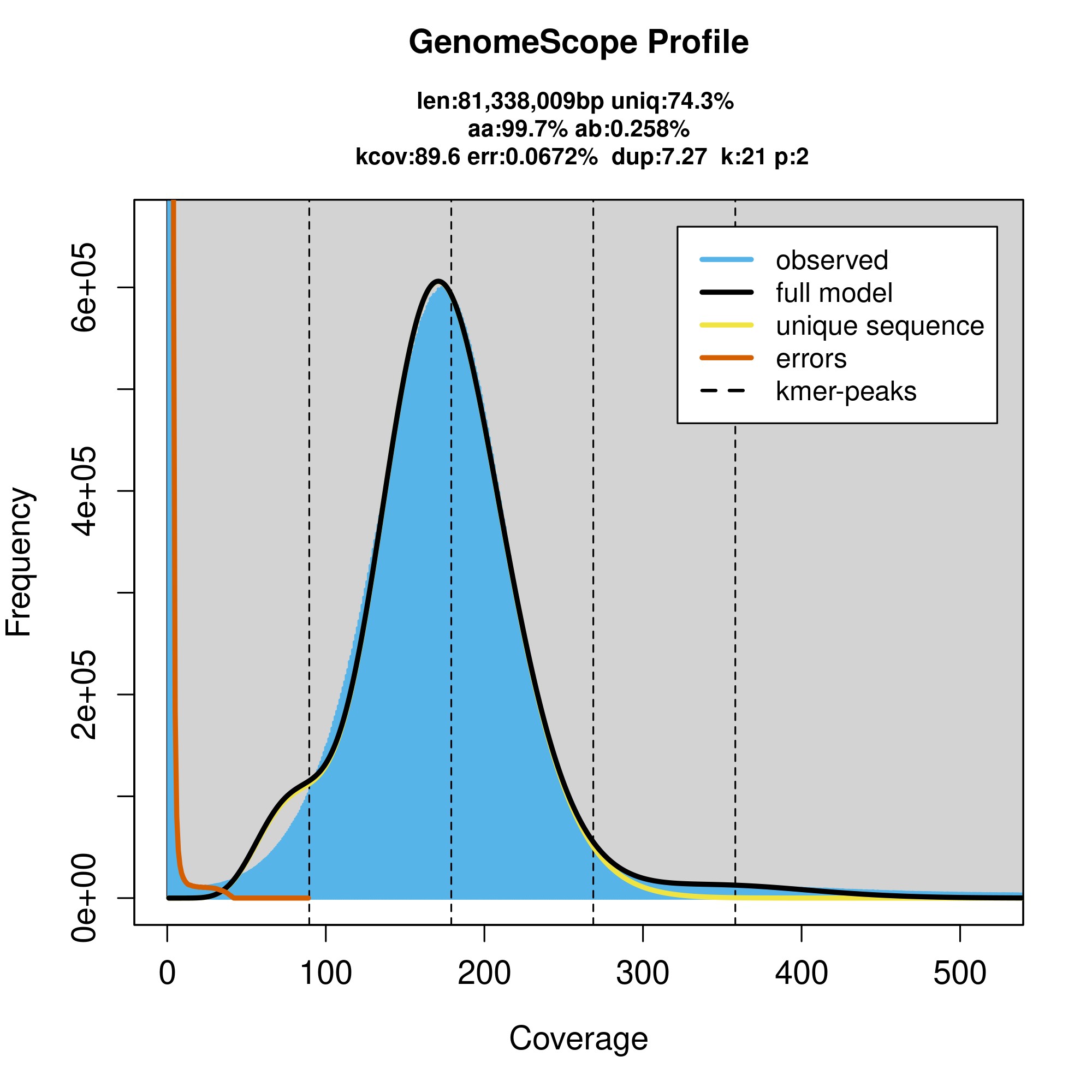

### Suppl. Figure 3.

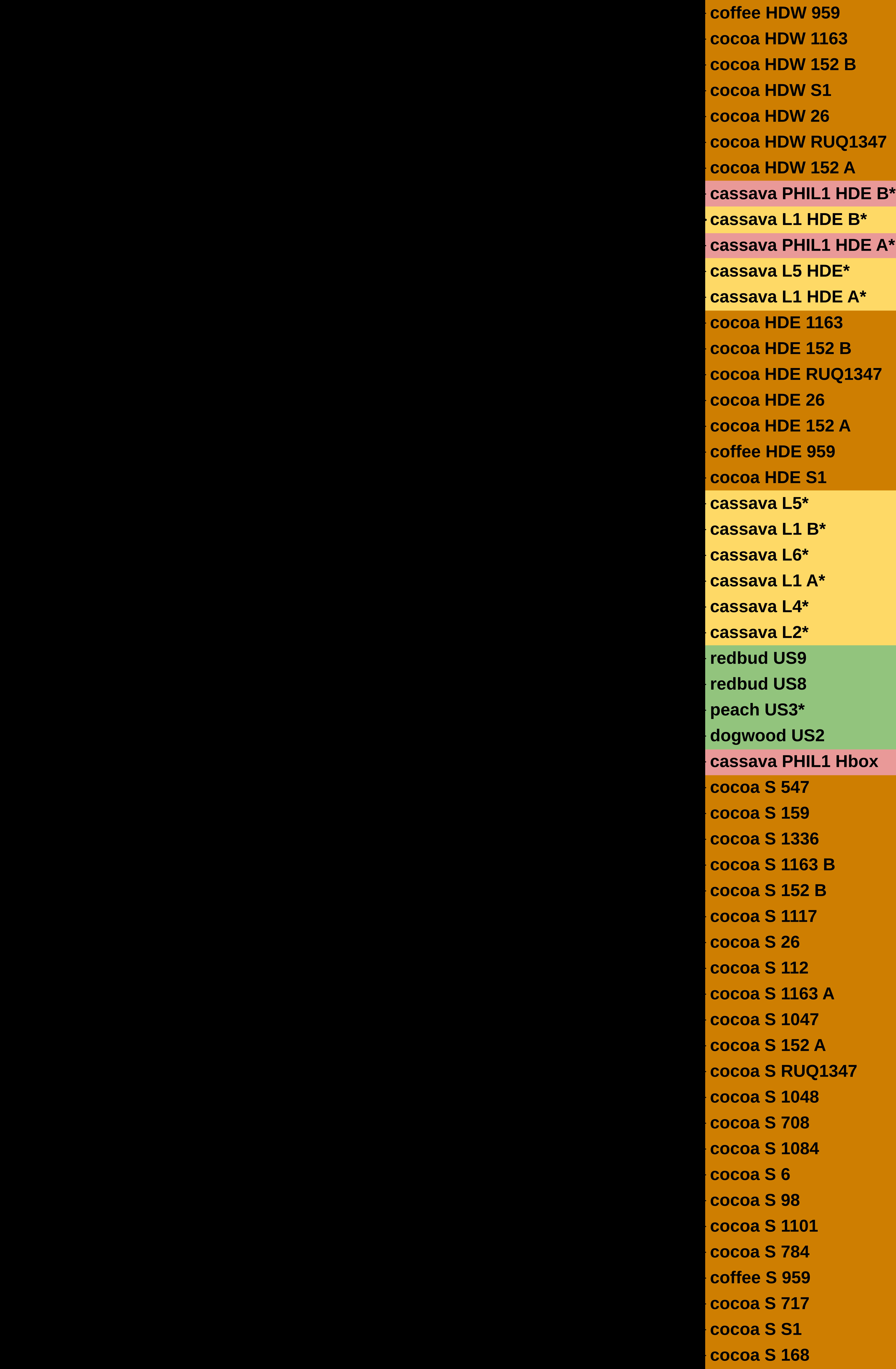

### Suppl. Figure 4.

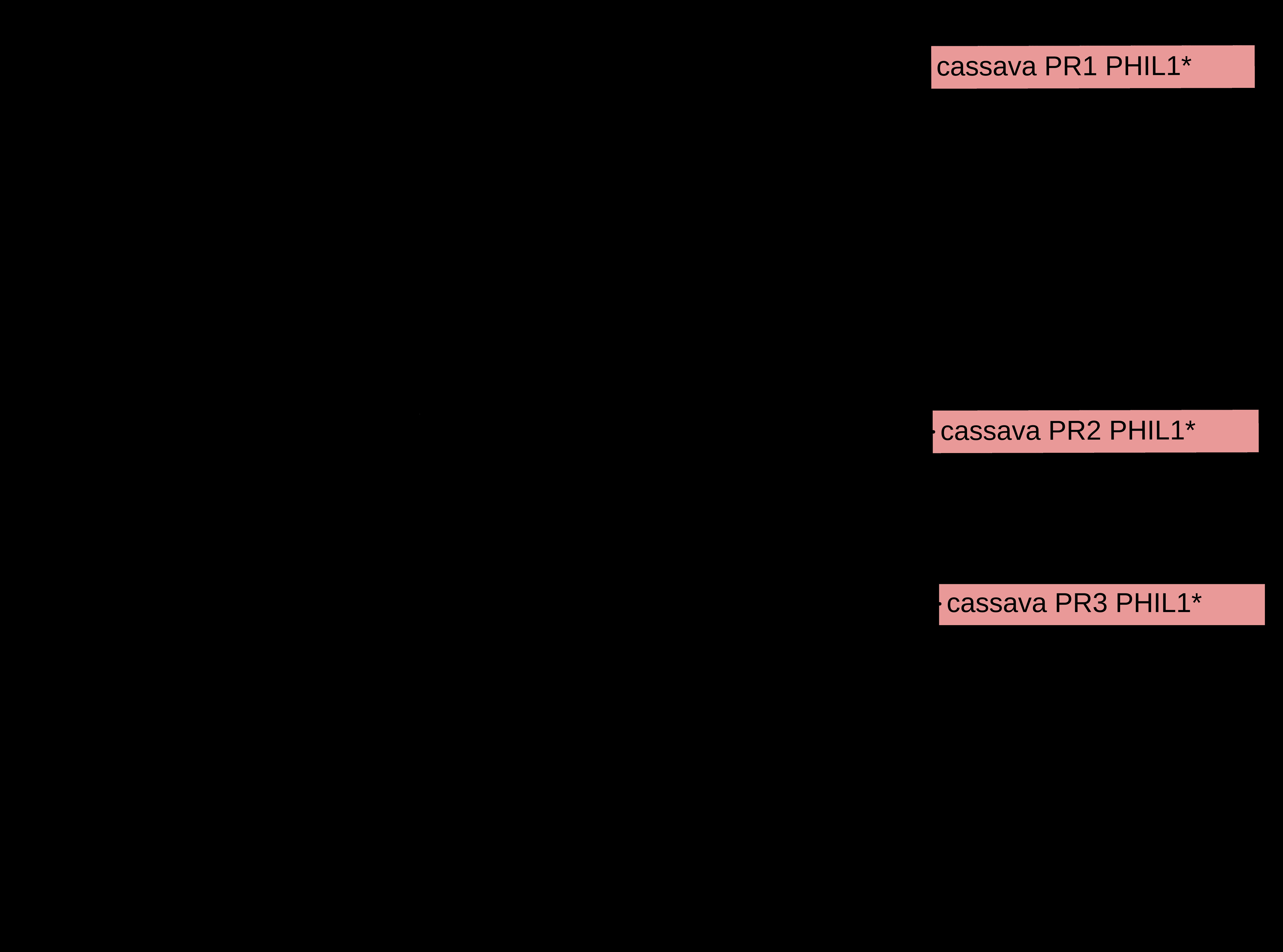
